# Cervicovaginal microbial features predict *Chlamydia trachomatis* spread to the upper genital tract of infected women

**DOI:** 10.1101/2024.11.26.625070

**Authors:** Sangmi Jeong, Tammy Tollison, Hayden Brochu, Hsuan Chou, Ian Huntress, Kacy S. Yount, Xiaojing Zheng, Toni Darville, Catherine M. O’Connell, Xinxia Peng

## Abstract

**Introduction:** *Chlamydia trachomatis (CT)* infection can lead to pelvic inflammatory disease, infertility and other reproductive sequelae when it ascends to the upper genital tract. Factors including chlamydial burden, co-infection with other sexually-transmitted bacterial pathogens and oral contraceptive use influence risk for upper genital tract spread. Cervicovaginal microbiome composition influences CT susceptibility and we investigated if it contributes to spread by analyzing amplicon sequence variants (ASVs) derived from the V4 region of 16S rRNA genes in vaginal samples collected from women at high risk for CT infection and for whom endometrial infection had been determined.

**Results:** Participants were classified as CT negative (CT−, n=77), CT positive at the cervix (Endo−, n=77), or CT positive at both cervix and endometrium (Endo+, n=66). Although we were unable to identify many significant differences between CT infected and uninfected women, differences in abundance of ASVs representing *Lactobacillus iners* and *L. crispatus* subspecies but not dominant lactobacilli were detected. Twelve informative ASVs predicted endometrial chlamydial infection (AUC=0.74), with CT ASV abundance emerging as a key predictor. We also observed a positive correlation between levels of cervically secreted cytokines previously associated with CT ascension and abundance of the informative ASVs.

**Conclusion:** Our findings suggest that vaginal microbial community members may influence chlamydial spread directly by nutrient limitation and/or disrupting endocervical epithelial integrity and indirectly by modulating pro-inflammatory signaling and/or homeostasis of adaptive immunity. Further investigation of these predictive microbial factors may lead to cervicovaginal microbiome biomarkers useful for identifying women at increased risk for disease.

## INTRODUCTION

*Chlamydia trachomatis* (CT) is the most common, sexually transmitted, bacterial infection globally (1), with young adults at greatest risk for infection. During childbirth, CT can also be transmitted to a newborn through contact with infected cervical tissue and secretions, resulting in infection of mucous membranes of the eye, oropharynx, urogenital tract, and rectum (2). In addition, CT has been extensively studied as a potential co-factor for human papillomavirus (HPV) enhancing cervical cancer risk through cellular transformation, viral load enhancement, and oncogene overexpression (3). When CT ascends from the cervix to the endometrium or uterus, it can lead to upper genital tract infection, increasing risk for pelvic inflammatory disease and reproductive sequelae such as ectopic pregnancy, or infertility (2). Despite the availability of diagnostics and effective treatments, the asymptomatic nature of CT infection often leads to undiagnosed cases, contributing to its importance as a global health burden (4).

The cervicovaginal microbiome (CVM) is frequently classified into community state types (CSTs), based on the dominant bacterial species present. Using 16s rRNA gene sequencing, Ravel et al. defined the most common CSTs (I-V) (5). CSTs I, II, III, and V are dominated by specific *Lactobacillus* species with overall lower diversity of other microbiota and are generally associated with healthy reproductive health outcomes (6, 7). However, CST IV is comprised of heterogeneous, predominantly anaerobic bacterial genera and *Lactobacillus* species are sparse (5). CST IV is associated with adverse health outcomes such as bacterial vaginosis (BV) (8), pregnancy failure (9), and pre-term birth (10). Vaginal dysbiosis, strongly associated with CST IV, has also been linked to increased STI vulnerability (7, 8, 11–13). In particular, *in vitro* (14, 15) and *in vivo* (16, 17) studies have associated indole-producers within the CVM with CT evasion of IFN-mediated depletion of tryptophan, an essential chlamydial nutrient (18, 19). Anaerobes also produce biogenic amines and proinflammatory short-chain fatty acids, promoting symptoms and disease (20, 21). In contrast, *Lactobacillus crispatus*-dominated CST I is considered protective against CT infection (8, 13, 22). Mechanisms contributing to protection include production of lactic acid that can directly kill CT (23) while D(-) lactic acid produced by some *Lactobacillus sp.* inhibits CT infection by down-regulating host epithelial cell proliferation (6). CVM-derived metabolites may also act on their host to stimulate DC activation and accelerate immunity (24). STI co-pathogens may also act directly, or in conjunction with microflora, to influence chlamydial infection. We previously observed that women with PID caused by *Neisseria gonorrhoeae* (NG) and CT carried systemic blood transcriptional signatures with greatest activation of cell death pathways and suppression of responses essential for adaptive immunity and CT infection clearance (25). Although these studies have established that CVM composition influences risk for chlamydial cervical infection (8, 13, 22, 26), the extent to which it influences ascending infection has yet to be investigated.

In this study, we sought to investigate the contribution of the CVM to ascending chlamydial infection using high-throughput amplicon sequencing based on 16S ribosomal RNA (16S rRNA) genes, by determining if bacterial members, especially those previously associated with susceptibility to cervical infection, can be implicated in chlamydial spread to the upper genital tract. We analyzed the amplicon sequence variants (ASVs) derived from the V4 region of 16S rRNA genes to investigate the CVM of study participants enrolled in T cell Response Against Chlamydia (TRAC), a highly characterized cohort of women at high risk for CT infection and for whom endometrial infection had been determined. When participants were categorized into CSTs based on their CVM composition, CST III (dominated by *Lactobacillus iners*; 30.0%) and CST IV (characterized as diverse; 54.5%) predominated. At the cervical level, ASV analysis revealed no significant overall differences in CVM composition between the CT−, Endo−, and Endo+ groups. Nevertheless, our analysis revealed that certain cervicovaginal microbial features were predictive of ascended chlamydial infection (Endo+). Including CT ASV abundance in the analysis significantly improved prediction accuracy, indicating that CT burden at the cervix is a critical factor in the progression of infection. Furthermore, we detected a positive correlation between microbial features predicting CT ascension and levels of selected cytokines in cervical secretions, indicating that these CVM members may influence ascending infection through their interaction with host immune responses, acting on CT indirectly and directly. This study underscores the potential of developing mucosal biomarkers to identify women at higher risk of CT upper genital tract infection, which could be valuable for clinical practice and future vaccine trials.

## RESULTS

### Classification of participants based on STI clinical diagnostic tests

Cervicovaginal swab samples and endometrial tissue biopsy specimens were collected from 246 women aged 18-35 years in the TRAC cohort (27, 28). Demographics and sexual history of the cohort were previously described (27). Cervical and endometrial infection status of study participants was determined using nucleic acid amplification tests (NAATs) on relevant samples (27). Twenty-six vaginal swab-derived samples were excluded from this study for the following reasons: three had quality concerns identified during DNA extraction, while the infection status of the remaining twenty-three samples was undetermined because they yielded equivocal results regarding chlamydial detection at the cervix or endometrium during clinical diagnostic testing.

The remaining 220 participants were classified into the following three groups: Endo+, Endo−, and CT− (**Supplementary Table S1**). A total of 143 women who tested positively for CT at their cervix (CT+) were subsequently divided into two groups based on confirmed upper genital tract infection. Women testing positively for CT infection at their cervix and endometrium (n=66) were defined as Endo− positive (Endo+) and the remaining CT infected women who tested positive for CT at their cervix but tested negative for endometrial CT infection (n=77) were grouped as Endo− negative (Endo−). The control group (n=77) was comprised of women testing negatively for cervical chlamydial infection (CT−). However, among the 77 uninfected women, seven samples from participants that were initially diagnosed as CT− by NAAT clinical testing but subsequently identified as CT+ by the detection of CT reads in 16S analysis, were later confirmed to be CT+ by separate quantitation of CT burden in cervical samples by qPCR (29). Consequently, these participants were excluded from 16S microbiome analyses comparing CT+ to uninfected, comparing Endo+ to Endo−, and were reserved for a proof-of-concept analysis. Clinical diagnostic testing at enrollment additionally determined cervical infection with the bacterial STI pathogens *Neisseria gonorrhoeae* (NG) and/or *Mycoplasma genitalium* (MG) in all participants (**Supplementary Table S1**) (27).

### Characteristics of cervicovaginal microbiota in the TRAC cohort

The CVM of TRAC participants were profiled using 16S rRNA gene V4 region sequencing and ASV-based analysis. ASVs identifying common STI bacterial pathogens were detected in cervicovaginal 16S rRNA including CT, NG, and MG (**Figure 1A**). CT ASV vaginal abundance amongst participants testing positively for CT infection positively correlated with CT cervical burden (Spearman correlation coefficient 0.575) (29) previously assayed by qPCR (27). CT ASV sequences were detected in 83 of the 143 CT− infected women (58%) and in 7 of 77 women who were diagnosed as CT− (9.1%) (**Figure 1A, Supplementary Table S1**). Of 20 NG infected women identified by clinical diagnostic testing, NG ASV sequences were detected in 17 of 20 (85%), while NG ASV sequences were detected in 5 out of 200 women testing negatively (2.5%). Assaying via qPCR indicated these discrepancies most likely were a consequence of low burden and/or differential site sampling (29). In contrast, MG-identifying ASV sequences were detected in only 6 of the 38 participants clinically diagnosed with MG (15.8%), an observation explained by the overall low burden of MG detected in infected TRAC participants (30). MG abundance in cervical specimens, previously determined by qPCR (30), was significantly associated with detection of MG ASV in sample libraries (Mann-Whitney, P=0.0206). Overall, these results support the high quality of this 16S rRNA based sequencing dataset.

**Figure 1.**
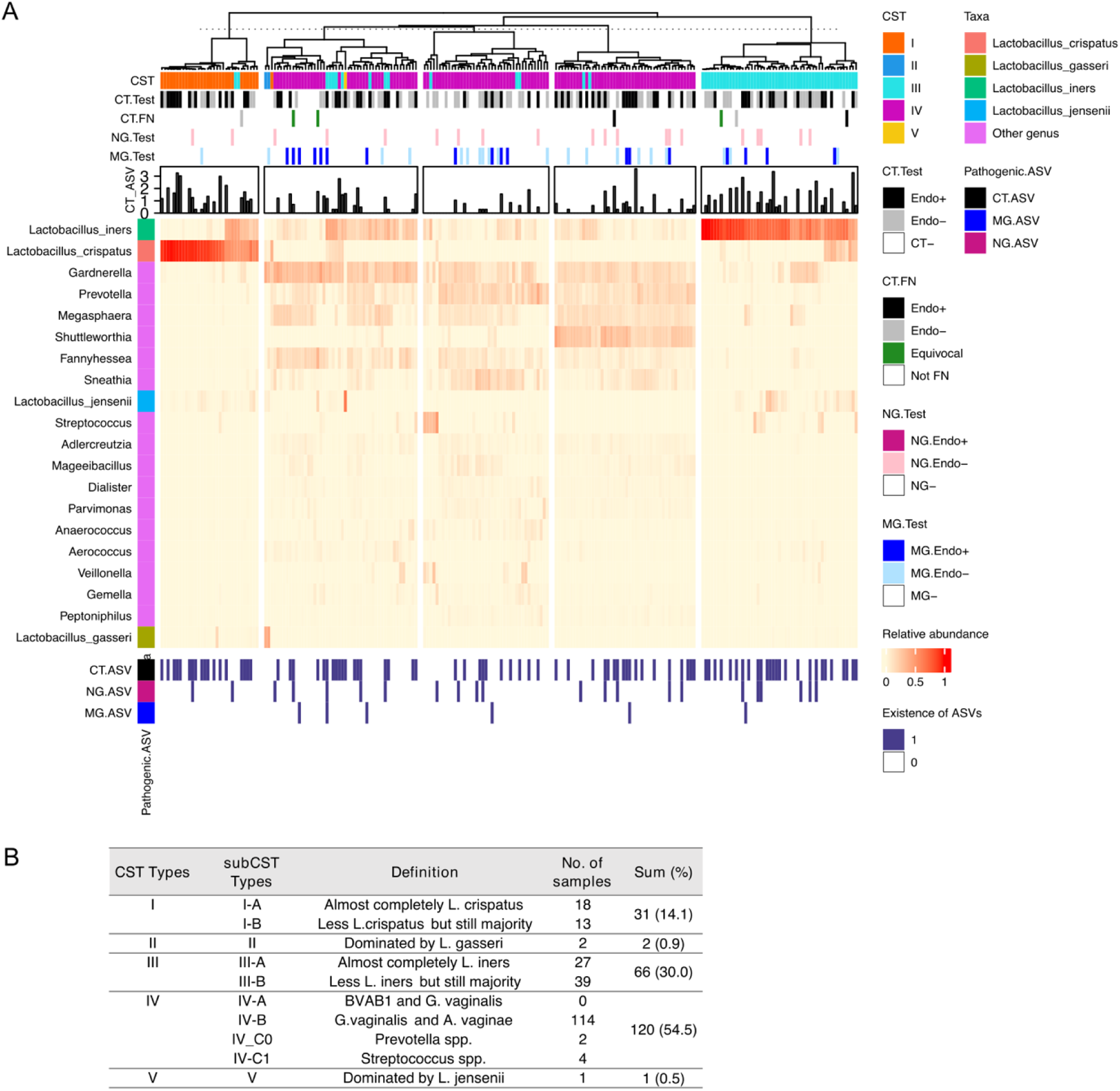
Profiles of community state types from TRAC cohort associated with Chlamydial infection. **A.** The taxonomic composition of 220 women, incorporating CST types and pathogen infection status (CT = *Chlamydia trachomatis*, NG = *Neisseria gonorrhoeae* (NG), and MG = *Mycoplasma genitalium*) were presented on the heat map. The testing results for pathogenic infection status at enrollment (CT.Test, NG.Test, MG.Test) were provided as bar graphs on the top of the heatmap. The reassigned results of seven false negatives for CT infection are labeled as CT.FN. The term ‘Equivocal’ indicates that the reassignment of these cases as either Endo+ or Endo− was unclear, despite them being confirmed as CT+. CT ASV read counts were converted into logarithmic scale (CT_ASV; log10(read count + 1)). The observations of three pathogenic ASV sequences were illustrated with ‘Existence of ASVs’ at the bottom of the heatmap (CT.ASV, NG.ASV, MG.ASV). **B.** The table shows the definitions of CST types and the number of women belonging to each CST type.

Based on ASV relative abundance from 213 women, the most prevalent genera in these women were *Lactobacillus, Gardnerella, Prevotella*, and *Megasphaera* (**Figure 1A, Supplementary Figure 1A**). The three most abundant ASVs observed overall were *Lactobacillus iners, Lactobacillus crispatus,* and *Gardnerella vaginalis* (**Supplementary Figure 1B**). Within each group (CT−, Endo−, and Endo+), individuals exhibited a spectrum of *Lactobacillus* abundance, from *Lactobacillus*-predominant compositions to *Lactobacillus*-sparse (**Figure 1A, Supplementary figure 1A-B**). Alpha diversity indices were assessed at the ASV level to investigate whether diversities of cervicovaginal microbial communities differed among the Endo+, Endo−, and CT− groups. Neither Shannon nor Inverse Simpson indices were significantly different between among three groups (**Supplementary Figure 1C**; Wilcoxon rank-sum test p-values > 0.05).

### Community State Types (CSTs) represented within TRAC cohort

The cervicovaginal microbial composition of TRAC participants was further evaluated by categorizing each sample according to CST, taxonomic profiles that represent discrete vaginal microbial communities (**Figure 1B**) (5, 31). The TRAC cohort exhibits a high prevalence of CT infection (65%, 143 of 220 participants) with CST IV (characterized as diverse) as the predominant CST type (54.5%, 120 of 220, **Figure 1**). Almost all the remaining participants were assigned as CST I (dominated by *L. crispatus;* 14.1%, 31 of 220) or CST III (dominated by *L. iners*; 30.0%, 66 of 220) (**Figure 1**). Two participants were assigned to CST II (dominated by *L. gasseri*) and one participant was assigned to CST V (dominated by *L. jensenii*) (**Figure 1**). The seven participants that tested false CT negative were assigned to CST I, III, and IV (**Supplementary Table S1**).

To assess a potential relationship between CST and epidemiological information in the TRAC cohort, we first evaluated the association between CST and cervical CT infection status for 213 women excluding seven false negatives (**Supplementary Table S1, Supplementary Table S2**). However, the association between CT infection status and CST was not significant (**Supplementary Table S2;** Fisher’s exact test P-value > 0.05). Second, the association between CST and demographics and physiologic information was assessed for 220 women (**Supplementary Table S1, Supplementary Table S2**). The CST type was significantly associated with Nugent score or race (**Supplementary Table S2;** Fisher’s exact test p-value < 0.05). Of the 119 women in CST IV, 107 (89.9%) were classified as having BV based on the Nugent score, compared to lower proportions of BV in the other CST types. CST IV was more common among Black and Multiracial participants, whereas CST I was more common among Caucasian participants, consistent with previous studies (5, 12, 32–35).

### Significant differences in abundance were observed between the CT+ and CT− groups for *Lactobacillus* and *Prevotella* species

Since it has been reported that cervicovaginal microbial profiles differ between CT− and CT+ individuals (36–40), we investigated CVM compositions of CT+ and CT− TRAC participants. We used sparse partial least squares discriminant analysis (sPLS-DA) (41) because this method is effective for supervised classification and feature selection, aiming to identify the best discriminating features between groups while handling high-dimensional data. The statistical discrimination performance for classifying CT− and CT+ women using the first two sPLS-DA components was estimated by a mean area under the curve (AUC) of 0.70 (**Figure 2A**), indicating that overall CVM profiles in the presence of CT ASV tended to differ between CT+ and CT− women. However, when CT ASV was excluded from the discrimination analysis, performance dropped to a mean AUC of 0.51 (**Figure 2B**).

**Figure 2.**
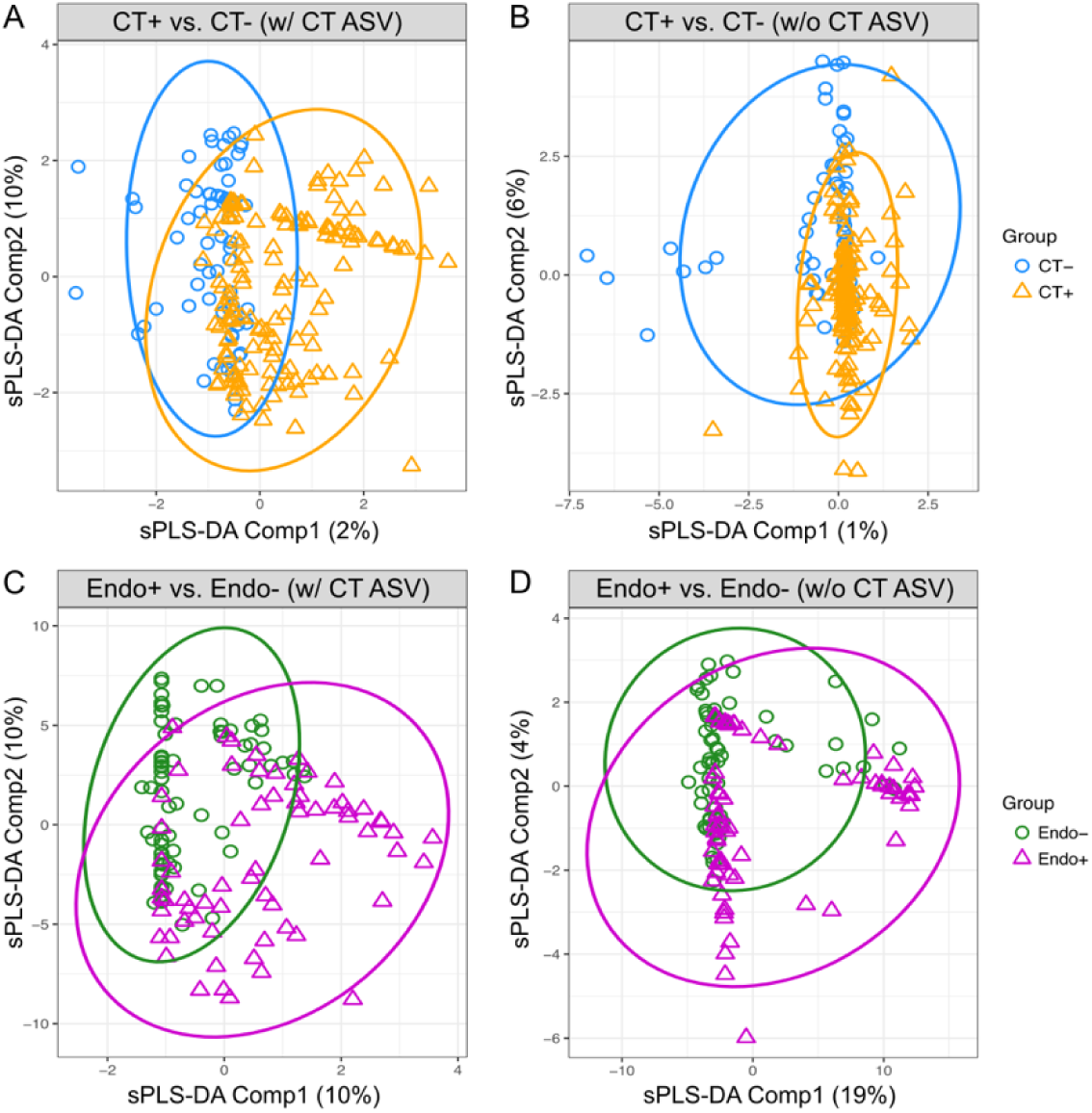
Discrimination of chlamydial infection status at the cervix o endometrium in TRAC cohort using sPLS-DA. Discrimination of chlamydial infection status at cervix (A and B) and endometrium (C and D) was visualized using the first two components of sPLS-DA. The discriminations were performed using ASVs including CT ASV (A and C) or excluding CT ASV (B and D). Each component’s contribution to the discrimination was indicated at their axis. The performance was evaluated using the mean Area Under the Curve (AUC) values. **A.** The first two sPLS-DA components discriminate CT+ from CT− women based on ASVs with CT ASV (AUC mean = 0.70, AUC s.d. = 0.02) **B.** The discrimination of CT+ from CT− women based on without CT ASV (AUC mean = 0.51, AUC s.d. = 0.03) **C.** The first two sPLS-DA components discriminate Endo+ from Endo− women (AUC mean = 0.76, AUC s.d. = 0.01) **D.** The discrimination of Endo+ from Endo− women based without CT ASV (AUC mean = 0.55, AUC s.d. = 0.03).

We also performed sPLS-DA analysis within CSTs, to examine if differences in CVM composition (excluding CT ASV) could be detected between CT+ and CT− individuals. In CST I (dominated by *L. crispatus*), CT− women appeared to cluster in a manner visually distinct from CT+ women but discrimination performance was statistically inconsistent, with a mean AUC of 0.49, likely due to the small number of samples assigned to CST I (n=30) (**Supplementary Figure 2A**). Within CST III (dominated by *L. iners*), cervical CT infection status was moderately discriminated (AUC mean = 0.71, AUC s.d. = 0.04, n = 63) (**Supplementary Figure 2B**) selecting six low-abundant *L. crispatus* ASVs for the first sPLS-DA component (**Supplementary Table S3**). Within CST IV-assigned participants, the discrimination between CT+ and CT− groups was weak (AUC mean = 0.50, AUC s.d. = 0.04, n = 117) (**Supplementary Figure 2C**).

To identify cervicovaginal microbiome features that are differentially abundant between CT− and CT+ groups, we used Microbiome Multivariate Association with Linear Models (MaAsLin2) (42), which produces a list of significant microbial features with false discovery rate correction. A total of 28 ASVs differed in abundance between CT+ and CT− groups, including ASVs assigned to *Fannyhessea vaginae*, *Gardnerella vaginalis* and multiple *Prevotella* spp, microorganisms previously associated with enhanced sensitivity to STI infection and BV (7, 8, 11–13) (**Supplementary Table S4**, unadjusted P-value < 0.05). CT (ASV sequence # ZOtu76) and NG (ASV sequence # ZOtu95) were more abundant in CT+ women, with Benjamini Hochbergh (BH)-adjusted P-values < 0.0001 and unadjusted P-values < 0.05, respectively. Two *Dialister* ASVs (ASV sequence #’s ZOtu105 and ZOtu220) were (**Supplementary Table S4**, unadjusted P-value < 0.05). Four of the 28 ASVs, including NG and 3 ASVs assigned to *Dialister invisus,* a member of the gut microbiome that has been associated with infections at other mucosal sites*, Prevotella sp.*, and *L. crispatus* (ASV sequence #’s Zotu96, Zotu220, Zotu304, and Zotu2052), were also identified as informative ASVs discriminating cervical CT infection status by sPLS-DA analysis (**Supplementary Table S3 and S4**).

Although no significant differences in abundance of dominant *Lactobacillus spp.* ASVs (ASV sequence #s ZOtu1, ZOtu2, ZOtu11, ZOtu32, and ZOtu374) were detected between CT+ and CT− participants in the overall cohort based on their predominant ASVs, differences were observed in 7 uncommon ASVs assigned to *L. iners*, *L. crispatus*, *L. jensenii*, *L. gasseri* and *L. colini* (**Supplementary Table S4, Supplementary Figure 3**). To investigate if these low-abundance ASVs were sequencing artifact, we conducted multiple sequence alignments to identify the position of mismatches between these low-abundant *Lactobacillus* ASV sequences and their corresponding reference sequence, reasoning that errors generated by merging of paired-end reads would be disproportionately concentrated in the middle of sequences. The 7 ASV sequences were aligned with the corresponding ASV sequences of their most abundant *Lactobacillus sp.* ASVs listed in **Supplementary Table S4**, and V4 region sequences of *Lactobacillus sp*. curated in the RDP 16S No18 reference database (43) (**Supplementary Figure 4**). The most abundant *Lactobacillus* ASVs (ASV sequence #s ZOtu1, ZOtu2, ZOtu11, ZOtu32, and ZOtu374) exactly matched their corresponding V4 region reference sequences (**Supplementary Figure 4**). In contrast, the 7 low-abundant *Lactobacillus* ASV sequences contained mismatches at 20 of 253 nucleotides in the V4 region, except for one ASV (ASV sequence # ZOtu459). Intriguingly, these nucleotide variants frequently occurred at the same 20 positions along the *Lactobacillus* V4 region, rather than being concentrated in the middle of ASV sequences (**Supplementary Figure 4**). These results suggested that the observed sequence variants in these low-abundant *Lactobacillus* ASVs reflected biological variation in *Lactobacillus* subspecies or strains that composed the CVM of TRAC participants.

We investigated whether CVM features could predict CT infection status within this high STI risk cohort, the majority of whom had BV at enrollment (27) (**Supplementary Table S1**). We performed a random forest-based analysis using ASV abundance, while excluding the CT ASV. In brief, we randomly partitioned the samples from CT+ and CT− groups into a training dataset and a separate test dataset. We built a predictive model using the training dataset and evaluated its performance by predicting the CT infection status using the independent test dataset (see Methods for details). The prediction performance was assessed using the average receiver operating characteristic (ROC) curves across 100 replicates. The corresponding average AUC was found to be 0.61 for the prediction using all ASVs excluding the CT ASV (**Supplementary Figure 5**). Thirteen ASVs contributed to prediction of absent CT infection (**Supplementary Table S5**). NG ASV (ASV sequence# ZOtu95), more abundant in CT+ women (**Supplementary Table S4**), was the strongest contributor to the prediction. NG is not considered a member of the CVM, thus its contribution to this prediction likely derives from the frequency of co-infection with CT in women reporting high risk behaviors for STI acquisition. Four *Syntrophococcus sucromutans-*assigned ASVs and four *Prevotella spp*. ASVs, which showed differential abundance between CT+ and CT− women (**Supplementary Table S4**), were found to be predictors of absent CT infection (**Supplementary Table S5**). However, the ASVs of *S. sucromutans* (ASV sequence #s ZOtu709, ZOtu1481, ZOtu1814, and ZOtu2372) have low confidence genus-level assignments, indicating that these strains, currently classified as members of the class *Clostridia*, require further investigation to resolve their taxonomic classification.

### Cervicovaginal microbial features are predictive of susceptibility to ascended chlamydial infection

Given the severe sequelae that can arise after CT infection ascends to involve the endometrium and fallopian tubes and the absence of diagnosing this outcome non-invasively, we investigated if CVM profiles of CT+ women were reflective of CT endometrial infection. We first examined the overall differences in CVM compositions between Endo− and Endo+ patients using sPLS-DA analysis, as before (see also Methods). Endo+ and Endo− women showed distinguishable separations by the first two sPLS-DA components including CT ASV (**Figure 2C**; AUC mean = 0.76, AUC s.d. = 0.01). The CT ASV (ASV sequence # ZOtu76) and two *Lactobacillus* ASVs (ASV sequence #s ZOtu915 and ZOtu914) contributed the most to the first component, while more than 50 ASVs were identified as important features for the second component discriminating Endo+ and Endo− women (**Supplementary Table S6**). Despite the high number of important features identified, in the absence of CT ASV, discrimination between Endo+ and Endo− weakened with an AUC of 0.55 (**Figure 2D**).

We also examined the CVM profiles within CST I, CST III, and CST IV types separately for CT endometrial infection status using sPLS-DA analysis. Endo− women were visually separable from Endo+ women within CST I with the first sPLS-DA component including 8 *Lactobacillus* ASVs (**Supplementary Table S6**), but the statistical performance was very low, with a mean AUC of 0.47 (**Supplementary Figure 2D**), again likely due to the small number of women within CST I (n=23). The discrimination of endometrial infection status within CST III was also weak (AUC mean = 0.58, AUC s.d. = 0.09, n=43) (**Supplementary Figure 2E**). However, endometrial infection status within CST IV (characterized as diverse) can be discriminated (AUC mean = 0.71, AUC s.d. = 0.03, n=76) (**Supplementary Figure 2F**) with the CT ASV solely contributing to the first sPLS-DA component and the interaction of 19 ASVs including 4 *Prevotella* ASVs and a *Mycoplasma microti*-assigned ASV (# ZOtu45) contributing the second sPLS-DA component (**Supplementary Table S6**).

To identify CVM features that would be most informative in predicting CT ascension, we performed random forest-based analysis using ASV abundances as features to predict individual patients in Endo+ or Endo− groups. Since CT is not only a key factor of ascended infection but potentially modulates the cervical and endometrial environment for other microbes, we assessed the predictive performance of the model in three scenarios: (A) prediction with all ASVs including CT ASV (# ZOtu76) (**Figure 3A**), (B) prediction with all ASVs but excluding CT ASV (**Figure 3B**), and (C) prediction with only CT ASV (**Figure 3C**). Average AUCs were 0.74 for the model with all ASVs, and 0.57 for the model that excludes the CT ASV (**Figure 3A-B**), indicating the importance of CT ASV abundance for prediction. When the model was trained by the CT ASV alone it resulted in an average AUC of 0.74 (**Figure 3C**).

**Figure 3.**
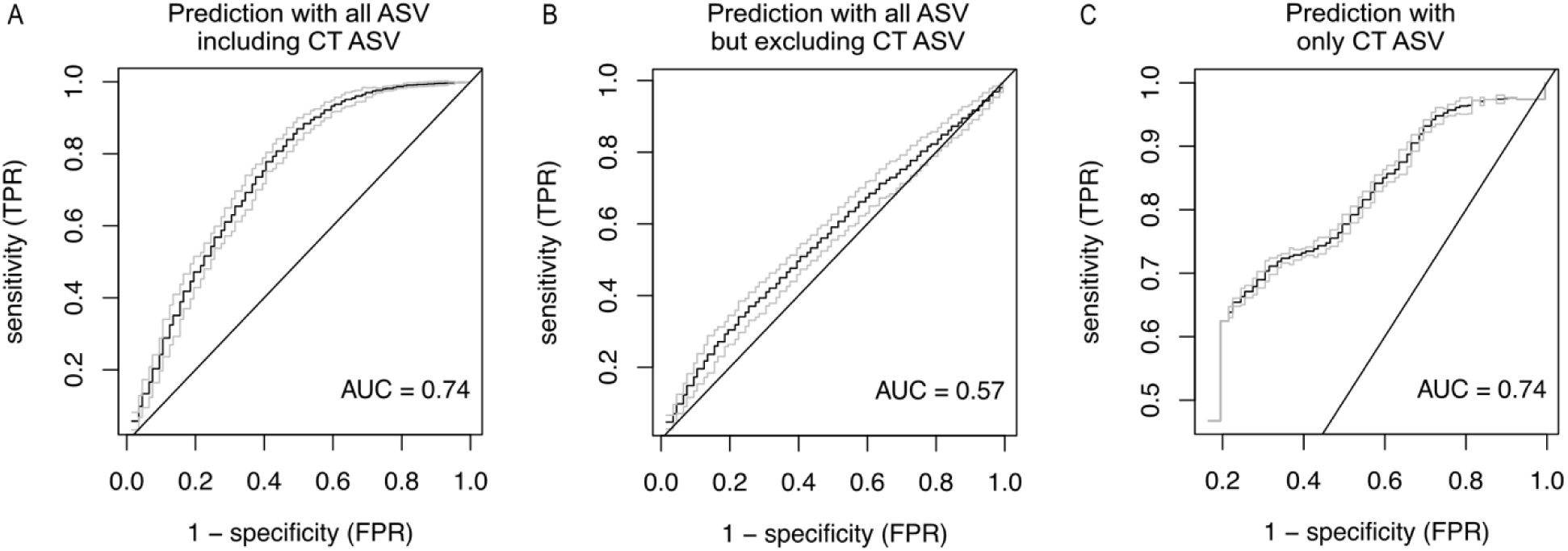
ROC curves and corresponding AUCs from random forest prediction for susceptibility to ascending chlamydial infection. Prediction accuracies were evaluated by averaging AUCs over 100 replicates for each scenario. **A.** The prediction performance for the susceptibility to CT ascension using all ASVs, including CT ASV resulted in an AUC of 0.74. **B.** The prediction performance for excluding CT ASV resulted in an AUC of 0.57. **C.** The susceptibility of ascending CT infection, predicted using only CT ASV, led to an AUC of 0.74.

In the prediction with all ASVs including CT ASV, CT and 11 additional ASVs were identified as informative features using selection frequency based on mean decrease in accuracy (**Table 1**; feature selection frequency 80 out of 100 replicates). These 11 ASVs were also selected as informative features in the prediction using all ASVs but excluding CT ASV (**Table 1**; feature selection frequency ≥ 80 out of 100 replicates). Notably, there was no overlap between these features and those selected as predictors of absent CT cervical infection (**Supplementary Table S5**). Among the identified 12 ASVs, four were identified with higher abundance in Endo+ women (ASV sequence #s ZOtu76, ZOtu233, ZOtu1808, ZOtu567) (**Table 1**). Of the four? highly abundant ASVs in Endo+ women, three were also identified as important features discriminating Endo+ and Endo− patients in the above sPLS-DA analysis (ASV sequence #s ZOtu76, ZOtu233, and ZOtu1808) (**Table 1** and **Supplementary Table S6**). All three ASVs assigned to *Sutterella stercoricanis* (ASV sequence #’s ZOtu110, ZOtu184, and ZOtu180) exhibited lower abundance in Endo+ women (**Table 1**). *Actinotignum schaalii* ASV (ASV sequence # Zotu281) and *Haemophilus haemolyticus* ASV (ASV sequence # Zotu103) were also identified as predictors, which were less abundant in Endo+. *Mycoplasma microti* (ASV sequence # Zotu45), a species previously isolated from the respiratory tracts of prairie voles (43), seemed an unlikely component of the CVM. Subsequent examination determined that the assigned ASV shared 100% identity with uncultured *Mycoplasma sp.* clone Mnola (44), now *Candidatus* Malacoplasma girerdii (45), a strict endosymbiont of *Trichomonas vaginalis* (TV). Within TRAC, we observed that detection of *Ca.* M. girerdii ASV reads coincided with a positive TV clinical diagnostic finding in 14 of 37 participants (37.8%) but was only detected in 7 of 183 (3.8%) of participants who tested negatively for TV suggesting that this was the more correct assignment. Overall, these results suggest that multiple CVM features together can be predictive of susceptibility to CT ascension but the amount of CT at the cervix is a key factor.

**Table 1.**
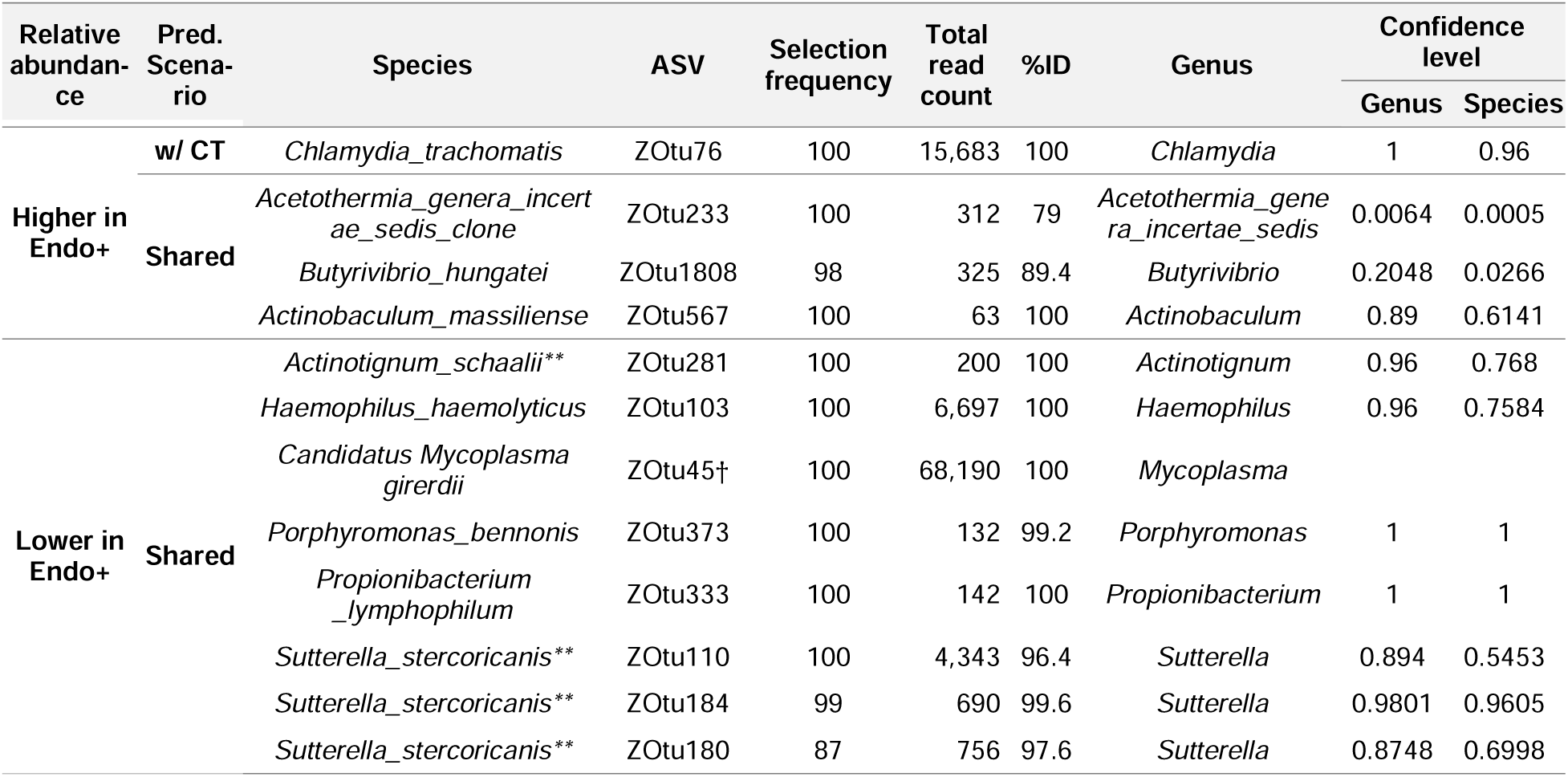
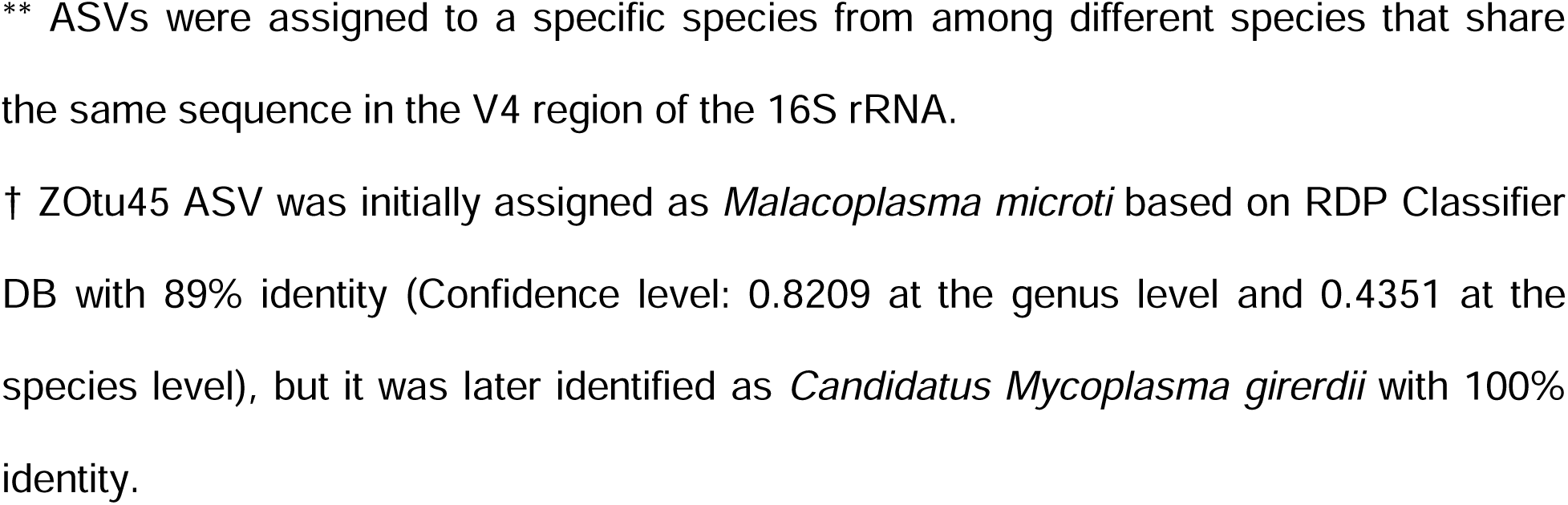
ASVs predictive of ascended CT infection. The ASVs selected for predicting susceptibility to ascending CT infection were determined based on their selection frequency threshold derived from random forest predictions. The selection frequency indicates the number of times an ASV was detected with non-NA values of mean decrease in accuracy, occurring in 9 or 10 out of 10 folds, out of 100 replications. From the RF classification results, 12 ASVs were identified as microbial biomarkers because their selection frequencies exceeded 80. Notably, despite considering two different scenarios (A: prediction with all ASVs including CT ASV and B: prediction with all ASVs but excluding CT ASV), 11 out of 12 ASVs were consistently identified across both scenarios and indicated by “Shared” in Prediction scenario column of Table 1. The relative abundance of these ASVs was determined using t statistics, indicating whether an ASV was higher in Endo+ or lower in Endo+. Total read count was calculated for all 220 women, and details of the %ID and Confidence level columns were provided in Table 1.

### Cervicovaginal microbial features associated with ascending CT infection are positively correlated with cytokines in cervical secretions associated with endometrial CT infection

To better understand possible functional consequences of these 12 CT ascension-related microbial features on host immune responses, we explored their associations with levels of 7 cytokines in TRAC cervical secretions that had been previously associated with CT ascension (44). The following cytokines, CXCL10, TNF-α, IL17A, CXCL9, CXCL11, CCL4, and CXCL13, were positively associated with endometrial infection (unadjusted P<0.05) in a univariable regression model. Using canonical correlation analysis (CCA) (45), we investigated whether there was an overall correlation between these cytokine levels and the predictive microbial features. With 12 ASVs, including CT, we observed a positive correlation between microbial abundances and cytokine levels (**Figure 4A**; canonical correlation = 0.54). There was also a positive but weaker association between cytokine levels and CVM-derived ASVs when CT ASV was excluded (**Figure 4B**; canonical correlation = 0.43). CT ASV abundance also correlated weakly with the seven cytokines (**Figure 4C**; canonical correlation = 0.49). Together, these results indicate that host immune responses are not exclusively modulated by CT abundance. The CCA-based analyses suggest that the predictive CVM microbial features influence CT ascension directly and indirectly, through interaction with host immune responses.

**Figure 4.**
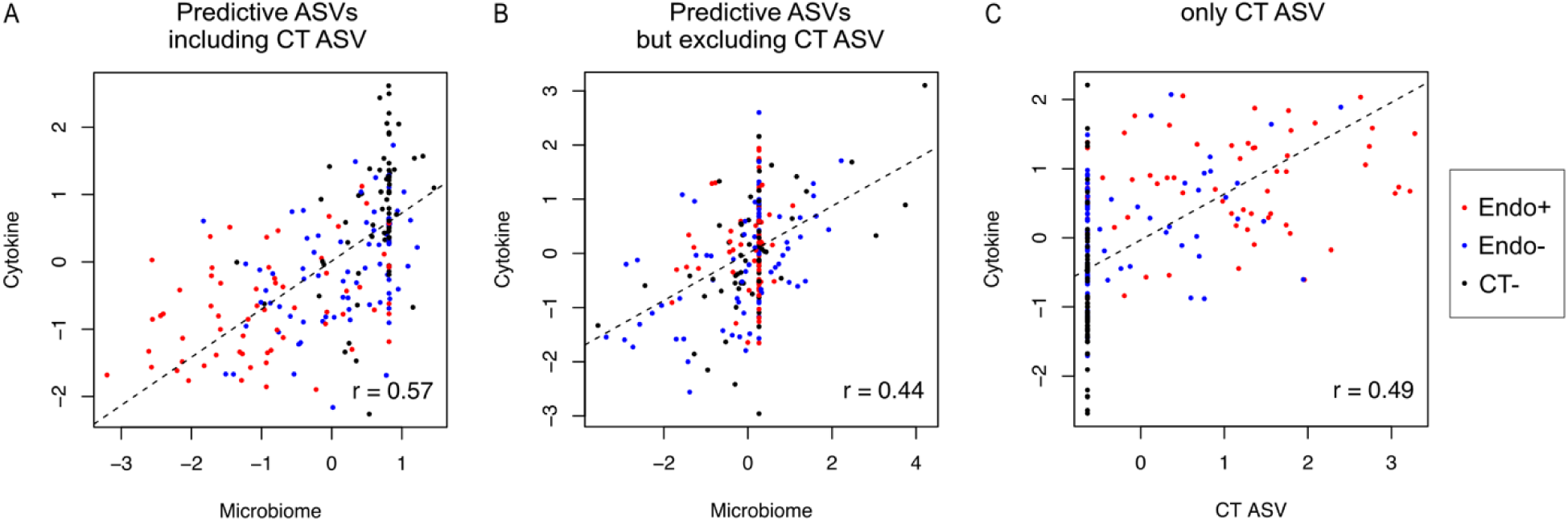
The Correlations between canonical variates of microbiome biomarkers and cervical cytokines. The canonical correlations between cervical cytokines related to chlamydial ascension and microbial biomarkers identified for chlamydial ascension risk were performed under two scenarios: (A) correlating the 7 cervical cytokine levels with 12 informative ASVs, including CT ASV, listed in **Table 1**, and (B) with 11 informative ASVs, excluding CT ASV. **A.** The canonical correlation between the seven cervical cytokine levels and the 12 ASVs, including CT ASV, was visualized using two CCA variates, resulting in a correlation coefficient of 0.57. **B.** The canonical correlation between the seven cervical cytokine levels and the 11 ASVs, excluding CT ASV, yielded a correlation coefficient of 0.44. **C.** Only CT ASV abundance was correlated with the seven cytokine levels, resulting in a correlation coefficient of 0.49.

### A proof-of-concept analysis for predicting ascending CT infection using the twelve informative ASVs

We identified twelve ASVs that are predictive of susceptibility to ascending CT infection using random forest-based prediction and ASV abundance of Endo+ and Endo− women (**Table 1 and Figure 3A**). Four false negative CT+ participants (**Supplementary Table S1**) had residual endometrium-derived samples for additional testing and were used for proof-of-concept prediction of Endo+ vs Endo− here. An optimal threshold of 0.55 was derived from the averaged ROC curve of the model informed by all ASVs (**Figure 3A**) and was used to determine between Endo+ and Endo− (**Supplementary Figure 6**). The model correctly predicted CT ascension status for two out of four participants based on confirmed detection of CT DNA (29). CT ASV abundance was considered as a key factor for the prediction of CT ascension but was unable to predict Endo+ vs Endo− status alone (data not shown). This proof-of-concept analysis of four false negative CT+ participants could not validate the 12 ASVs including CT ASV due to very small sample size. Validation of the 12 CVM features identified requires testing in a larger, independent cohort.

## DISCUSSION

Recent literature has revealed that cervicovaginal microbiota may modulate susceptibility to infections such as human papillomavirus (HPV), CT, NG and MG infections (46). CVM composition has been associated with increased risk of incident CT infection (16, 22, 46, 47). In this study, we sought to determine whether CVM composition facilitates CT ascension from the cervix to the upper genital tract by profiling 16S rRNA genes obtained from vaginal swabs collected at enrollment from the TRAC cohort. Although we were unable to identify many significant differences between CT infected and uninfected women, we identified 12 cervicovaginal microbial features that predicted active upper tract infection. Chlamydial cervical burden was a key driver of ascent because vaginal CT ASV abundance alone, effectively predicted this outcome with AUC of 0.74 for sensitivity and specificity. We also observed positive correlation between levels of 7 cytokines previously associated with CT ascension and abundance of CVM-derived ASVs, indicating that local host immune responses during infection were not exclusively modulated by pathogen load.

TRAC is comprised of women at elevated risk for STI infection and a striking feature of this cohort is the very high rate of CT infection, 67%, amongst enrollees, reflective of highly targeted recruitment strategies (27). This may have blunted our ability to detect significant differences in the composition of the CVM, at the level of CST, for individual genera or species that associated with protection against CT infection. Unsurprisingly, most participants were assigned to CST III or CST IV, previously recognized as neutral or favorable for CT infection (7, 8, 11–13). Within CST III, CT− women were separable from CT+ using sPLS-DA components containing *L. crispatus*-assigned ASVs. Amongst CST I-assigned participants, the CST most highly associated with protection from CT infection (8, 13, 22), CVM composition did not differ sufficiently to cluster CT infected and uninfected participants effectively. TRAC was designed as a longitudinal study to investigate T cell responses important for protection from incident chlamydial infection, but the high rate of CT positivity at enrollment combined with a stringent requirement that participants have completed a minimum of three follow up assessments, provided insufficient power to confirm previously identified associations of CST with infection susceptibility. Nevertheless, a random forest-based analysis using all ASVs (excluding CT) was moderately predictive of CT− women **(Supplementary Table S5)**, while MaAsLin2 analysis identified differentially abundant community members previously associated with susceptibility **(Supplementary Table S4)**. Conserved between both analyses were ASVs assigned to *Prevotella sp.* and *S. sucromutans* that associated with CT infection. Studies, *in vitro* (14, 15) and *in vivo* (16, 17) have associated indole-producers, e.g. *Prevotella sp*., with CT evasion of IFNγ-mediated tryptophan depletion (18, 19). Of the *Prevotella sp*. discriminating CT infection in our analysis (48), only *P. brunnea* is an indole producer (48, 49), but it is also possible that they modulate host factors, indirectly influencing chlamydial susceptibility. *S. sucromutans*, in association with *Prevotella sp*., serves as a biomarker of gut barrier dysfunction (leaky gut) (50) where increased luminal abundance of Zonulin-1, an endogenous tight-junction protein, was associated with decreased abundance of these species. Directly relevant to our study, recent work by Hinderfield and colleagues (51), demonstrated that CST IV-associated microorganisms promote paracellular permeability of human ectocervical monolayers with coincident down-regulation of Zonulin and Occludin expression, leading them to hypothesize that undermining cervical epithelium integrity enhances susceptibility to TV and other STI pathogens (51). Thus, a CVM that contributes essential substrates and improves pathogen access to the cervical epithelium can increase susceptibility to chlamydial infection.

In contrast, lactic acid produced by *Lactobacillus sp.* directly kills CT (23) while D(-) lactic acid produced by some *Lactobacillus sp.* inhibits CT infection by down-regulating host epithelial cell proliferation (6). In this study, we were unable to detect associations of the most abundant *Lactobacillus* species with CT positivity but identified differences in low abundance strains **(Supplementary Table S4)**. We questioned whether the low-abundant *Lactobacillus* ASVs that were differentially abundant by MaAsLin2 analysis were genuine *Lactobacillus* species or accidental assignments of artificial sequences to *Lactobacillus* species, given the taxonomic complexity among *Lactobacillus* species (52) and the limitations of characterizing these species through 16S rRNA gene sequencing (53). Although ASVs have intrinsic biological meaning independent of reference database or the specific context of a study (54), the taxonomic resolution is much lower compared to the full length of 16S rRNA sequences or metagenome sequences (55). Recent genomic studies identifying genetic elements that control functional differences between and within *Lactobacillus species*, such as the role of bile salt hydrolases that promote colonization of vertebrate-adapted niches (56), and the ability to metabolize extracellular glycogen important for microbial growth in the female genital tract (57, 58) as well as identification of novel urogenital tract species (59) suggest that it is premature to conclude that the differences we detected between CT infected and uninfected women are artifactual, and future investigations should consider implementing typing approaches with superior resolution that will enable us to determine if further investigation into the biological roles of individual *Lactobacillus* subspecies in cervical CT infection is needed.

The TRAC cohort provided a unique opportunity to investigate the CVM contribution to chlamydial ascension from the cervix to the upper genital tract. Using sPLS-DA, we were able to observe overall differences in CVM compositions between Endo− and Endo+ participants, although discriminatory resolution dropped considerably when the CT ASV was excluded, emphasizing the importance of CT cervical abundance in driving ascension. At the level of CST, discriminatory performance was highest in CST IV, but much reduced for CST I and CST III participants, suggesting that CVM-derived factors that promote susceptibility to CT infection also contribute to its ascent. However, we did not detect much overlap in the specific CVM features differentially abundant between CT− and CT+ participants by MaAsLin2 analysis and those selected by random forest analysis as predictive of spread with the obvious exception of the CT ASV.

Many of the CVM features selected by our analysis have not previously been associated with human chlamydial infection but have been investigated in the context of the respiratory or gut microbiome. Through “nutritional immunity”, the host may limit iron availability as a means of protection against invading pathogens (60). CT requires iron (61, 62) and may experience additional competition for this essential nutrient from CVM members. Iron restriction also impacts host cellular metabolism, leading to differential host gene expression with downstream consequences for availability of other key substrates required for chlamydial developmental success (63). Antithetically, transcription of the *trpBA* operon essential for chlamydial scavenging of exogenous indole to tryptophan is iron-regulated (64) leading to the proposal that a low iron environment could increase risk for sequelae (64). *H. haemolyticus* is a respiratory tract commensal that limits infection with the pathogenic *H. influenzae* through competition (65), because it uses its superior ability to sequester iron to facilitate occupation of their shared niche (66, 67). Its relatively lower abundance in Endo+ participants may indicate that chlamydial need for iron predominates during natural infection and that *H. haemolyticus* abundance modulates ascended infection through its negative influence on chlamydial growth.

Also less abundant in Endo+ participants, *S. stercoricanis* has not been previously described as a significant member of CVM but within the human gut microbiome, its reduced abundance has been identified as a predictor of enhanced susceptibility to respiratory infection (68). *Sutterella sp.* are epithelium-adherent (69), but do not contribute to gut dysbiosis or loss of gut epithelial integrity. In the context of STI infection, their abundance is significantly increased in the rectum of HIV negative, STI uninfected men, and CD4/CD8 ratio positively correlated with *Sutterella* in HIV-infected patients, consistent with an association with T cell-mediated adaptive responses independent of active bacterial infection (70). They are mildly pro-inflammatory, and it has been proposed that their association with their human host is mutualistic through their contribution to steady state induction of Th17 cell differentiation (71, 72). We previously reported that IL-17A transcript was significantly increased in blood of participants with high cervical *C. trachomatis* burden compared to those with low cervical burden and uninfected participants (25) and that cervical secretion of IL-17A was associated with increased odds of endometrial infection (44). Nevertheless, we observed increased frequencies of CD4 Th17 cells in peripheral blood of CT− infected TRAC participants compared to uninfected participants, and a further increase in participants without recurring infection compared to those who were reinfected with CT within a year after enrollment (73). Together, these results suggest that CT infection elicits local and systemic Th17 responses, and that the systemic Th17 responses detected were associated with reduced reinfection. It is possible that occupation of a niche by benign *S. stercoricanis* within the CVM of Endo− participants protects against expansion of members that positively contribute to chlamydial burden by provision of necessary intermediates or by compromise of epithelial integrity. It is also possible that its reduced abundance in Endo+ women contributed to reduced CD4 T cell activity, ultimately delaying resolution of chlamydial infection. Future studies that are more informative of function, e.g. transcriptomics, metabolomics, may identify common metabolites or pathways that can be associated with CT spread to the upper reproductive tract, while profiling of adaptive immune cell populations recruited to the infection site will lend further insight into mechanisms of chlamydial clearance.

We previously observed the importance of CT/NG coinfection on risk for CT reinfection (27). We also reported altered bloodborne transcriptional signatures pointing to suppressed host adaptive responses with CT/NG coinfection in women with pelvic inflammatory disease (25). However, we did not observe TV nor MG co-infection associating with ascended CT in TRAC participants (27, 30). In this study, an ASV that we ultimately re-assigned to *Ca*. M. girerdii (74, 75) was identified as a CVM feature predictive of ascended infection. Members of this large and diverse class of *Mollicutes,* distinguished by a small genome and lacking a cell wall, such as *M. hominis* and *Ureaplasma sp.*, have been identified as commensals and/or opportunistic genital tract pathogens, although meta-analyses have not effectively resolved the extent of their impact on female infertility or other reproductive outcomes including preterm birth (76, 77). Intriguingly, *M. hominis* and *Ca. M. girerdii*, alone or together, can live intracellularly or on the surface of TV, accelerating the parasite’s growth (78, 79) and modulating its pro-inflammatory potential. Relevant to CT infection because of the importance of cell-mediated responses for chlamydial clearance, TV harboring *M. hominis*, but not alone, synergistically upregulates IL-8, IL-1β, and TNF-α production as well as inducing production of the Th-17 polarizing cytokine IL-23 by human monocytic THP-1 cells with coincubation (80). If, as we speculate, recruitment of chlamydia-specific Th1 and Th17 cells to the genital tract is essential for control, TV coinfection, in some but not all instances, might promote this outcome, explaining why *Ca.* M. girerdii ASV was associated with Endo− women. Confusingly, ASVs assigned to the other TV endosymbiont *M. hominis* were also detected in many TRAC participants but were not selected as predictive features. We were not able to immediately resolve this contradictory observation because rRNA from TV was not captured in this analysis since it is a protozoan and the clinical diagnostic was not quantitative, but it highlights the potential for factors that influence pathogen success to be mediated through interactions between microbiome members, including those that cannot be detected using a 16S rRNA profiling approach.

ASVs contributing to prediction of ascended CT infection that were more abundant in Endo+ women included *Actinobaculum massiliense,* a urinary tract commensal or opportunistic pathogen (81), and reportedly a rare cause of PID (82). A recent proteomic analysis identified two highly expressed subtilisin-like proteases which may cleave cytokines (83), providing potential to modulate the host inflammatory response to itself and CT. ASVs assigned to *Butyrivibrio hungatei* and the genus *Acetothermia* were also more abundant in Endo+ participants. Neither bacterium is well documented as a member of CVM. In the context of gut microflora, *B. hungatei* is thought to act positively because reduced abundance of this butyrate producer has been associated with dysbiosis and disorders of the gut-brain axis such as Alzheimer’s Disease (84). Butyrate is a short chain fatty acid (SCFA) that modulates colon health by multiple mechanisms. In low amounts it serves as a carbon source for other commensals, while at higher doses it can be toxic, targeting other members of normal flora or pathogens such as *Salmonella sp*. (85). Colonic enterocytes transport butyrate as a preferred substrate (86) promoting epithelial health and integrity. However, in the female genital tract, SCFAs are associated with the production of pro-inflammatory cytokines and perturbation of epithelial integrity (87).

With potential translational impact, random-forest analysis revealed the importance of cervical CT abundance in prediction of concomitant upper genital tract spread. Using the CT ASV alone is a highly informative and simple feature, capable of predicting chlamydial ascension with equivalent AUC performance to the prediction using 12 ASV CVM biomarkers. A major barrier to preventing CT disease is delayed diagnosis, because infection is often asymptomatic (88), but both clinical and subclinical upper-tract inflammation can lead to chronic oviduct damage (89). Diagnostic biomarkers of ascended infection would be particularly useful for evaluation of therapeutics and vaccines (90–92) and improve case management, since women with clinical PID receive a longer antibiotic course compared to asymptomatic women with CT (93). Abundance of the CT ASV appears to be a better predictor of ascended CT infection than blood-borne transcriptional biosignatures (94) or profiling circulating immune cells (73) for this cohort of highly exposed women but was unsuccessful when applied to our small proof-of-concept set. Nevertheless, CT ASV with the CVM-derived selective features provided the best correlation with local cytokines, indicating the potential for further improved prediction with inclusion of host immune features. We previously performed a small pilot study that indicated that whole transcriptome sequencing analysis of ribosomal RNA-depleted total RNA isolated from cervical swab samples contained pathogen-specific sequences from women with confirmed sexually transmitted bacterial pathogens (95). Simultaneously, we identified and quantified their active microbial communities and after integration with their associated host-derived reads, we detected clustered host transcriptional profiles reflecting microbiome differences and STI infection. Together, these findings indicate that cervical sampling can yield sensitive and robust biomarkers for evaluation of candidate chlamydial vaccines, such as those currently in development or advancing towards Phase I (96) and Phase II trials.

High throughput sequencing of 16S rRNA enabled us to efficiently identify CVM communities in the TRAC cohort, but taxonomic resolution for microbial species or strains was frequently too low to confidently determine their taxonomy as cervicovaginal microbiome. Nonetheless, we were able to detect STI pathogenic microbes, as well as highly variable microbes such as *Lactobacillus sp.* and *Mycoplasma sp.* using ASV sequences. Copy number of the 16S rRNA gene can vary widely, ranging from 1 to 37 in bacteria and 1 to 5 in archaea (97). This variation in 16S rRNA gene copy number (GCN) limits the accuracy of quantifying 16S rRNA genes and CT 16S reads, potentially introducing biases in subsequent analyses (98). Indeed, this effect may be reflected in our variable success detecting ASVs associated with NG (four copies) versus CT (two copies) and MG (single copy) (99) in samples from women testing positively with highly sensitive clinical diagnostic tests. Although novel methods have been developed to correct for 16S rRNA GCN (100–103), we did not apply 16S rRNA GCN prediction tools for precise quantification of 16S rRNA genes in this study. In future work, we can adopt full-length 16S rRNA sequencing as an alternative to short reads of 16S rRNA to improve taxonomic resolution and quantification accuracy.

This study had several additional limitations. For example, the participants of TRAC cohort were highly exposed to CT infection, which may have introduced bias in the representation of the CVM communities in general. Vaginal swab samples for microbiome sequencing were collected only at enrollment when participants were actively infected, so there were no baseline samples to investigate temporal changes in CVM features related to the development of CT ascending infection. Even if we had chosen to sample participants longitudinally, all received antibiotic treatment at enrollment, which could have affected CVM composition. Also, 16S rRNA-based sequencing provided information only on the composition of CVM profiles. This limits the ability to understand the functional roles that these microbial communities may play in modulating host immune resistance or susceptibility to CT ascension. We did not investigate other factors such as host genetic heterogeneity and the effect of innate immunity on ascended CT infection.

In future studies, the integration of other techniques such as transcriptomics and metabolomics with 16S rRNA based sequencing, studies that have been initiated with establishment of an additional cohort of CT− infected women (TRAC2) (104) are expected to facilitate the development of deeper functional insights and identification of relevant biological pathways. Additionally, expanding the analysis with other relevant members of the CVM including parasites such as TV and viruses could also enhance our understanding of microbial interactions with the host and improve the prediction of upper genital tract infection spread. These multi-omics analyses may also contribute to the identification of biomarkers of protection or disease risk that could contribute to assessment of vaccine efficacy in future clinical trials.

## ACKNOWLEDGEMENTS

We thank the women who agreed to participate in this study; Ingrid Macio, Melinda Petrina, Carol Priest, Abi Jett, and Lorna Rabe under direction of Harold Wiesenfeld and Sharon Hillier, for their efforts in the clinic and the microbiology laboratory; and the staff at the Allegheny County Health Department STD Clinic, for their efforts. We would also like to acknowledge the contribution of Dr. Ian Huntress, who sadly passed away during the performance of these studies.

## Availability of data and material

Raw 16S rRNA gene sequencing data supporting the conclusions of this article is publicly available at NCBI’s Short Read Archive (SRA) under BioProject accession number PRJNA1136868 (https://www.ncbi.nlm.nih.gov/bioproject/). **Supplementary Figures S1–S8** and **Supplementary Tables S1–S3, S5**, and **S7** are included in **Additional File 1: Supplementary Materials**. **Supplementary Table S4** is provided in **Additional File 2**, and **Supplementary Table S6** is provided in **Additional File 3.** The legends of **Supplementary Tables S4** and **S6** are provided in **Additional File 1: Supplementary Materials.**

## METHODS

### T Cell Response Against Chlamydia (TRAC) Cohort

This cohort was composed of mainly young (median age, 21 years; range, 18-35 years), single (89%), African American (66%) women at high risk for the acquisition of CT (**Supplementary Table S1**). Inclusion criteria indicating high-risk status included > 3 sexual partners in the previous 6 months, ≤ 14 years of age at sexual debut, history of pelvic inflammatory disease (PID), or presentation to the recruitment site with any of the following: presence of mucopurulent cervicitis on exam, or sexual contact with a partner known to be infected with CT or NG or non-gonococcal non-chlamydial urethritis. Women with a current diagnosis of PID according to the Centers for Disease Control and Prevention guidelines were excluded. Additional exclusion criteria were pregnancy, uterine procedure or miscarriage in the preceding 60 days, menopause, hysterectomy, antibiotic therapy in the preceding 14 days, and allergy to study medications. A total of 246 TRAC participants were enrolled into a longitudinal study designed to investigate T cell responses important for protection from incident chlamydial infection over 12 months of follow-up between February 2011 and August 2014 and were recruited from the Allegheny County Health Department’s Sexually Transmitted Diseases Clinic, Magee-Womens Hospital (MWH) Ambulatory Care Clinic, and the Reproductive Infectious Disease Research Unit at MWH in Pittsburgh, PA. This study complied with the Declaration of Helsinki guidelines and all study participants provided informed consent prior to initiation of study procedures. The Institutional Review Boards at the University of Pittsburgh and the University of North Carolina approved the study. Participants completed questionnaires regarding obstetric/gynecologic history, behavioral practices, sexual histories, contraceptive methods, and symptoms according to study protocols. Clinical, histological, and microbiological testing was performed, and blood and endometrial samples were obtained at enrollment, after which all participants received single-dose agents to treat *gonorrhoea* (ceftriaxone, 250 mg intramuscularly) and chlamydia (azithromycin, 1 g orally). Participants in this cohort were assessed for cervical and endometrial infection using Aptima Combo 2 (AC2) (Hologic, Marlborough, MA) NAAT with overall infection rates of 67% and 8.5% for CT and NG, respectively. MG and TV infection was determined using Aptima MG and Aptima TV diagnostics (Hologic) respectively.

### 16S library preparation and sequencing

DNA was extracted from cervicovaginal swabs stored at −80 ℃ using the AllPrep DNA/RNA Mini Kit (Qiagen, Valencia, CA). Before extraction, the manufacturer’s protocol was optimized with minor modifications to achieve the best performance in terms of DNA/RNA yield and equal recovery of six different species comprising the ATCC Vaginal Microbiome Whole Cell Mix (MSA-2007). Average yields were 52.8 ng/µl yield of DNA in a final volume of 100µl per sample. The V4 regions of 16S rRNA gene were amplified using the Illumina 16S V4 primer set of 515F [GTGYCAGCMGCCGCGGTAA] and 806R [GGACTACNVGGGTWTCTAAT] (105, 106) designed for dual indexing as described by the Earth Microbiome Project protocol (https://earthmicrobiome.org/protocols-and-standards/16s/). Sequencing libraries were prepared according to the Illumina’s 16S Metagenomic Sequencing Library Preparation Guide (https://support.illumina.com/downloads/16s_metagenomic_sequencing_library_preparation.html; Part # 15044223 Rev. B), and sequenced on an Illumina NextSeq 500 Sequencer (Illumina, San Diego, CA).

### 16S rRNA-seq data processing

The 16S rRNA data analysis pipeline is outlined in **Supplementary Figure 7**. Raw fastq files containing paired-end read sequences per each sample were demultiplexed and preprocessed into amplicon sequence variants (ASVs) using bioinformatics software, USEARCH v11.0 (107) and PEAR (108). The fastq_eestats2 function in USEARCH was used to determine if trimming was needed to optimize read merging, based on expected errors for 150bp forward and reverse reads across different length cutoffs (100, 110, 120, 130, 140, and 150bp), which was 150bp for our data. Merged reads were expected to be >252bp with short overlap nucleotides (<10bp). PEAR, known for its robustness with short overlapping regions, was employed for read merging, utilizing a minimum overlap size of 8 bases (108). Following read merging, primer sequences were trimmed from both ends, and low-quality reads were filtered at a maximum expected error rate of 1.0.

Sequence denoising (error-correcting) and the generation of zero-radius OTUs (zOTUs), referred to as amplicon sequence variants (ASVs), were achieved using unoise3 function in USEARCH (107). The unoise3 algorithm prioritizes read abundance over sequence differences by processing sequences based on descending order of read abundance (109). As a result, a rare sequence can be incorporated into a highly abundant centroid sequence even if its difference is relatively high and may be discarded by chance depending on the order of pooled unique sequences (107, 109). To reproduce the same set of ASV sequences in TRAC cohort despite this limitation of randomly discarding low-abundant sequences, we added additional steps to the standard denoising and ASV generation process. We reordered unique sequences based on read frequency, read length, entropy, and confidence level of taxonomy assignment, all in decreasing order except for read length. Considering the optimal length of V4 region of 16S rRNA DNA to be 253bp, we calculated the distances between 253bp and the length of read, and then arranged the reads from the smallest distance to the longest distance away from 253bp. Using the rearranged ∼995,000 unique read sequences, we consistently reproduced the same total set of 2,777 ASVs from an average of ∼151,000 reads per sample. Of 2,777 ASV sequences, the Burrows-Wheeler Alignment tool (110) aligned 569 to the human reference genome (GRCh38) and these were excluded. We also removed ASV sequences shorter than 252bp, with 1,990 ASVs retained for further analysis. The inferred ASVs were assigned taxonomy using the RDP Classifier 16S trainset No. 18 raw training database (https://sourceforge.net/projects/rdp-classifier/files/RDP_Classifier_TrainingData/).

For analyses based on ASV abundances of CT+ and CT− women, 783 out of 1,990 ASVs were excluded because they had fewer than 50 read counts across CT+ and CT− groups because these low-abundance ASVs represented experimental artifacts generated during sequencing. Similarly, of 1,990 ASVs, 1,043 with fewer than 50 reads across Endo+ and Endo− women were excluded from analyses based on ASV abundances of Endo+ and Endo− women.

Multiple sequence alignment of 7 low-abundant *Lactobacillus sp*. ASV sequences (**Supplementary Table S4**) with their corresponding sequences of the most dominant *Lactobacillus sp*. ASVs and the V4 region sequences of *Lactobacillus sp*. obtained from the RDP 16S No18 reference database was performed using MUSCLE 3.8.31v (111). The alignment was visualized using UGENE (112) to with the goal of identifying mismatches in the central region of the 252bp ASV sequences (**Figure S4**).

### Modification of reference sequences for V4 regions in RDP Classifier

RDP Classifier 16S trainset No18 raw training database provides both bacterial and archaeal 16S rRNA reference sequences. Among a total of 21,195 reference sequences, nearly 40% had identical V4 region sequences for multiple species ranging from 2 to 65 different species in the RDP Classifier database (**Supplementary Table S7**). In such cases, sintax function for taxonomic assignment in USEARCH randomly chooses a species among multiple species given by the reference annotation (113). In addition, since the RDP Classifier database is not limited to human-related microbial taxonomy annotation, assigned species may include those not typically observed in the human cervicovaginal microbiota. For example, although *L. crispatus* and *L. jensenii* are commonly found in the cervicovaginal environment, taxonomic assignment based on the V4 region of the RDP Classifier database might incorrectly identify *L. acidophilus* as the predominant *Lactobacillus* species instead (114). To enhance the reliability of species assignments, we implemented a two-step approach. From the ASV abundance table, we selected a single species among those with identical V4 reference sequences based on the highest abundant species driven from the RDP Classifier database. Selected species are indicated with stars (**) in **Tables 1** and **Supplementary Table S4**. Remaining species and their corresponding V4 reference sequences were removed from the reference database. Using this reduced set of V4 region sequences as our reference database, we repeated taxonomy assignment for ASVs. Despite random, sintax algorithm-driven selection of a species from among multiple alternatives, we were able to track the species chosen from among those with identical V4 sequences.

### Statistical analyses

#### Alpha diversity

To investigate significant differences in alpha diversity among the three groups (Endo+, Endo−, and CT−), we standardized the total read counts per sample, which ranged from 34,571 to 417,336, by rarefying the reads across samples to the minimum read counts (34,571 reads) using rarefy_even_depth function in phyloseq R package. We assessed Shannon (richness) and Inverse Simpson (evenness) using phyloseq and microbiome R packages. Phylogenetic diversity was calculated using picante R package. The phylogenetic tree for phylogenetic diversity was generated from the 16S rRNA V4 region sequences of swab samples by using MUSCLE (111). Statistical significance in alpha diversity between each pair of groups was determined using the Wilcoxon rank-sum test.

#### The centered log-ratio transformation

Microbiome-derived read count data obtained through amplicon sequencing is analyzed as compositional data (115) where a proportional change for any microorganism will affect relevant abundances of others, rendering ASV read abundances mutually dependent. Centered log-ratio (CLR) transformation is one approach that generalizes relative abundances with respect to the geometric mean of all sequences in the sample (116). CLR-transformation of read counts enables meaningful comparisons of microbial abundances by mitigating compositional data biases (117). Since CLR-transformation depends on logarithms, handling zero counts is challenging. We treated absolute zero counts by replacing them with the smallest CLR-transformed values calculated by from our read counts.

#### Community state type (CST) classification of TRAC cohort and discrimination analysis for CT infection status associated with CST

TRAC participants (n=220) were categorized into CST types according to their vaginal microbial compositions using VALENCIA (VAginaL community state typE Nearest CentroId clAssifier) (118). VALENCIA classifies samples for CST types based on their similarity to a set of reference centroids, and the VALENCIA tool and reference centroids were sourced from GitHub (https://github.com/ravel-lab/VALENCIA). Where necessary, the taxon names from RDP Classifier database were manually adapted to correspond with the VALENCIA taxonomy of the reference centroids.

A sparse partial least squares discriminant analysis (sPLS-DA) (41) was employed to discriminate between CT infection status across all women, and to differentiate between CT infection status within CST I, CST III, and CST IV types. The sPLS-DA was executed using splsda function in mixOmics R package (119). For training across 213 women, 10-fold cross-validation (CV) with 1,000 iterations was used, while within assigned CSTs, sPLS-DA utilized 5-fold CV repeated 1,000 times. To assess the sPLS-DA model, AUC scores were computed from the training CV sets and averaged over 1,000 iterations, yielding the mean and standard deviation (s.d.). The ASVs listed in **Supplementary Table S3** and **Supplementary Table S4** were extracted using the perf function in the mixOmics R package with a threshold of frequencies greater than 0.80 for each component.

#### Microbiome Multivariate Association with Linear Models (MaAsLin2) for differential abundance analysis

To investigate if CVM of TRAC cohort differ in abundance between CT+ and CT− women, MaAsLin2 (42) was performed using Maaslin2 R package with the embedded CLR transformation method (normalization = “CLR”, transform = “NONE”, analysis_method = “LM”, correction = “BH”, min_prevalence = 0).

#### Random forest classification

Random forest classification was used to determine if CVM excluding CT is predictive of the lack of CT infection. This method was also utilized to predict chlamydial ascending infection and to identify microbial biomarkers. The random forest was performed using randomForest R package (120), and the processes of parameter selection, and biomarker identification, are outlined in **Supplementary Figure 8**. In the parameter selection, we selected the optimal hyperparameters to improve prediction performance. For model evaluation and biomarker identification, a random forest classification model was generated and evaluated with K-fold CV, simultaneously choosing the microbial biomarkers of chlamydial ascension.

The following parameters were examined: (1) number of folds for CV (K=5 or K=10), (2) level of phylogeny used to aggregate ASVs into species or genus, (3) significance level of T-tests and Wilcoxon rank-sum tests used for aggregated microbial features (0.05 or 0.10), (4) whether to use the default setting of randomForest function or a hyperparameter grid search. Tuning parameters in the randomForest function included three options: mtry (number of trees randomly sampled as candidates at each split), nodesize (minimum size of terminal nodes), and sampsize (sizes of sample to grow). To accomplish this, we split our dataset into training and test datasets using K-fold CV, ensuring that each fold included samples from both the Endo+ and Endo− groups. With training data, we performed two-sample T-tests and Wilcoxon rank-sum tests on ASVs to aggregate ASVs having p-value < 0.05 and the same direction in abundances within the same genus or species between Endo+ and Endo− groups. Specifically, when ASVs have a p-value less < 0.05, ASVs demonstrating the same directional trend in abundances (i.e., either positive or negative t-statistics) within a species or genus were aggregated into a single microbiome feature. ASVs with p-values >0.05 but with consistent directional trends, were also aggregated into a single microbiome feature. These aggregated microbiome features were tested again using T-tests and Wilcoxon-tests to choose microbial features having p-values less than the significance level (0.05 or 0.10). The aggregated microbiome features were subsequently employed as features of training data and test data for random forest classification. For every combination of the four parameters, AUC values were calculated from 20 replicates of random forest, and parameters were selected based on the highest sum of the mean and median AUC values across the 20 replicates. As a result, the optimal parameters chosen included a significance level of 0.10, a 10-fold cross-validation (CV), aggregation of read counts at the genus level for ASVs, and the default settings of the randomForest R function. These settings were applied twice, once including CT ASV as a microbiome feature, then excluding CT ASV.

With optimal parameters, the process re-implemented for model evaluation and biomarker identification with 100 iterations. Prediction accuracy was assessed by averaging the ROCs and AUCs obtained from the test sets over the 100 iterations. Optimal thresholds for specificity and sensitivity were obtained by pROC R package (121). For biomarker identification, ASVs were ranked based on their mean decrease in accuracy, which indicates the importance of each feature in classifying samples. The higher the mean decrease in accuracy, the more important the variable is in the model. Using this information, ASVs were identified as biomarkers predicting chlamydial ascension.

For validation of the 12 informative ASVs indicating risk for chlamydial ascension, we performed random forest prediction using the same approach by which the 12 informative ASVs were identified (**Table 1**). A total of 143 samples (both Endo+ and Endo−) were randomly split into training and test datasets using 10-fold cross-validation (CV). The test dataset, including the four samples, was predicted using the CLR-transformed abundance of the 12 ASVs. This entire process was repeated 100 times to ensure consistency in the predictions across different subsets of the 143 samples used as the training dataset. We then evaluated the predictions by calculating the mean, standard deviation, and standard error of Endo+ probabilities for the seven samples, as well as for the Endo+ and Endo− samples.

#### Canonical Correlation Analysis

Canonical correlation analysis (CCA) focuses on finding linear combinations maximizing the correlation between two sets of multiple variables. To assess the association between cervical cytokine levels and microbial features, CCA was implemented by CCA R package (45).

## Bibliography

1. Cheong HC, Lee CYQ, Cheok YY, Tan GMY, Looi CY, Wong WF. Chlamydiaceae: Diseases in Primary Hosts and Zoonosis. Microorganisms. 2019;7(5).

2. O’Connell CM, Ferone ME. Chlamydia trachomatis Genital Infections. Microb Cell. 2016;3(9):390–403.

3. Silva J, Cerqueira F, Medeiros R. Chlamydia trachomatis infection: implications for HPV status and cervical cancer. Arch Gynecol Obstet. 2014;289(4):715–23.

4. Paavonen J. Chlamydia trachomatis infections of the female genital tract: state of the art. Ann Med. 2012;44(1):18–28.

5. Ravel J, Gajer P, Abdo Z, Schneider GM, Koenig SS, McCulle SL, et al. Vaginal microbiome of reproductive-age women. Proc Natl Acad Sci U S A. 2011;108 Suppl 1(Suppl 1):4680–7.

6. Edwards VL, Smith SB, McComb EJ, Tamarelle J, Ma B, Humphrys MS, et al. The Cervicovaginal Microbiota-Host Interaction Modulates Chlamydia trachomatis Infection. mBio. 2019;10(4).

7. Borgdorff H, Tsivtsivadze E, Verhelst R, Marzorati M, Jurriaans S, Ndayisaba GF, et al. Lactobacillus-dominated cervicovaginal microbiota associated with reduced HIV/STI prevalence and genital HIV viral load in African women. ISME J. 2014;8(9):1781–93.

8. van Houdt R, Ma B, Bruisten SM, Speksnijder A, Ravel J, de Vries HJC. Lactobacillus iners-dominated vaginal microbiota is associated with increased susceptibility to Chlamydia trachomatis infection in Dutch women: a case-control study. Sex Transm Infect. 2018;94(2):117–23.

9. Tsai HW, Tsui KH, Chiu YC, Wang LC. Adverse effect of lactobacilli-depauperate cervicovaginal microbiota on pregnancy outcomes in women undergoing frozen-thawed embryo transfer. Reprod Med Biol. 2023;22(1):e12495.

10. Tabatabaei N, Eren AM, Barreiro LB, Yotova V, Dumaine A, Allard C, et al. Vaginal microbiome in early pregnancy and subsequent risk of spontaneous preterm birth: a case-control study. BJOG. 2019;126(3):349–58.

11. Gosmann C, Handley SA, Farcasanu M, Abu-Ali G, Bowman BA, Padavattan N, et al. Lactobacillus-deficient cervicovaginal bacterial communities are associated with increased HIV acquisition in young South African women. Immunity. 2017;46(1):29–37.

12. Anahtar MN, Byrne EH, Doherty KE, Bowman BA, Yamamoto HS, Soumillon M, et al. Cervicovaginal bacteria are a major modulator of host inflammatory responses in the female genital tract. Immunity. 2015;42(5):965–76.

13. van der Veer C, Bruisten SM, van der Helm JJ, de Vries HJ, van Houdt R. The Cervicovaginal Microbiota in Women Notified for Chlamydia trachomatis Infection: A Case-Control Study at the Sexually Transmitted Infection Outpatient Clinic in Amsterdam, The Netherlands. Clin Infect Dis. 2017;64(1):24–31.

14. Ziklo N, Huston WM, Taing K, Katouli M, Timms P. In vitro rescue of genital strains of Chlamydia trachomatis from interferon-gamma and tryptophan depletion with indole-positive, but not indole-negative Prevotella spp. BMC Microbiol. 2016;16(1):286.

15. Beatty WL, Belanger TA, Desai AA, Morrison RP, Byrne GI. Tryptophan depletion as a mechanism of gamma interferon-mediated chlamydial persistence. Infection and immunity. 1994;62(9):3705–11.

16. Ziklo N, Vidgen ME, Taing K, Huston WM, Timms P. Dysbiosis of the Vaginal Microbiota and Higher Vaginal Kynurenine/Tryptophan Ratio Reveals an Association with Chlamydia trachomatis Genital Infections. Front Cell Infect Microbiol. 2018;8:1.

17. Jordan SJ, Olson KM, Barnes S, Wilson LS, Berryhill TF, Bakshi R, et al. Lower Levels of Cervicovaginal Tryptophan Are Associated With Natural Clearance of Chlamydia in Women. J Infect Dis. 2017;215(12):1888–92.

18. Ziklo N, Huston WM, Hocking JS, Timms P. Chlamydia trachomatis Genital Tract Infections: When Host Immune Response and the Microbiome Collide. Trends Microbiol. 2016;24(9):750–65.

19. Aiyar A, Quayle AJ, Buckner LR, Sherchand SP, Chang TL, Zea AH, et al. Influence of the tryptophan-indole-IFNgamma axis on human genital Chlamydia trachomatis infection: role of vaginal co-infections. Front Cell Infect Microbiol. 2014;4:72.

20. Yeoman CJ, Thomas SM, Miller ME, Ulanov AV, Torralba M, Lucas S, et al. A multi-omic systems-based approach reveals metabolic markers of bacterial vaginosis and insight into the disease. PLoS One. 2013;8(2):e56111.

21. Ness RB, Kip KE, Hillier SL, Soper DE, Stamm CA, Sweet RL, et al. A cluster analysis of bacterial vaginosis-associated microflora and pelvic inflammatory disease. Am J Epidemiol. 2005;162(6):585–90.

22. Parolin C, Foschi C, Laghi L, Zhu C, Banzola N, Gaspari V, et al. Insights Into Vaginal Bacterial Communities and Metabolic Profiles of Chlamydia trachomatis Infection: Positioning Between Eubiosis and Dysbiosis. Front Microbiol. 2018;9:600.

23. Gong Z, Luna Y, Yu P, Fan H. Lactobacilli inactivate Chlamydia trachomatis through lactic acid but not H2O2. PLoS One. 2014;9(9):e107758.

24. Pulendran B, Maddur MS. Innate Immune Sensing and Response to Influenza. Curr Top Microbiol Immunol. 2014.

25. Zheng X, O’Connell CM, Zhong W, Nagarajan UM, Tripathy M, Lee DA, et al. Discovery of blood transcriptional endotypes in women with pelvic inflammatory disease. The Journal of Immunology. 2018;200(8):2941–56.

26. Filardo S, Di Pietro M, Porpora MG, Recine N, Farcomeni A, Latino MA, et al. Diversity of Cervical Microbiota in Asymptomatic Chlamydia trachomatis Genital Infection: A Pilot Study. Front Cell Infect Microbiol. 2017;7:321.

27. Russell AN, Zheng X, O’Connell CM, Taylor BD, Wiesenfeld HC, Hillier SL, et al. Analysis of Factors Driving Incident and Ascending Infection and the Role of Serum Antibody in Chlamydia trachomatis Genital Tract Infection. J Infect Dis. 2016;213(4):523–31.

28. Russell AN, Zheng X, O’Connell CM, Wiesenfeld HC, Hillier SL, Taylor BD, et al. Identification of Chlamydia trachomatis Antigens Recognized by T Cells From Highly Exposed Women Who Limit or Resist Genital Tract Infection. J Infect Dis. 2016;214(12):1884–92.

29. Jeong S, Tollison T, Brochu H, Chou H, Yu T, Baghaie P, et al. Aptima Combo2-avoiding variants detected in cervical and endometrial specimens from a cohort of sexually active cisgender women during 16S microbiome profiling. medRxiv. 2024.

30. Taylor BD, Zheng X, O’Connell CM, Wiesenfeld HC, Hillier SL, Darville T. Risk factors for Mycoplasma genitalium endometritis and incident infection: a secondary data analysis of the T cell Response Against Chlamydia (TRAC) Study. Sex Transm Infect. 2018;94(6):411–3.

31. Zhou X, Brown CJ, Abdo Z, Davis C, Hansmann MA, Joyce P, et al. Disparity in the vaginal microbial community composition of healthy Caucasian and black woman. ISME J. 2007;1:121–33.

32. Fettweis JM, Brooks JP, Serrano MG, Sheth NU, Girerd PH, Edwards DJ, et al. Differences in vaginal microbiome in African American women versus women of European ancestry. Microbiology (Reading). 2014;160(Pt 10):2272–82.

33. Vodstrcil LA, Twin J, Garland SM, Fairley CK, Hocking JS, Law MG, et al. The influence of sexual activity on the vaginal microbiota and Gardnerella vaginalis clade diversity in young women. PLoS One. 2017;12(2):e0171856.

34. Borgdorff H, van der Veer C, van Houdt R, Alberts CJ, de Vries HJ, Bruisten SM, et al. The association between ethnicity and vaginal microbiota composition in Amsterdam, the Netherlands. PLoS One. 2017;12(7):e0181135.

35. Brotman RM, He X, Gajer P, Fadrosh D, Sharma E, Mongodin EF, et al. Association between cigarette smoking and the vaginal microbiota: a pilot study. BMC Infectious diseases. 2014;14(471):1–11.

36. Ceccarani C, Foschi C, Parolin C, D’Antuono A, Gaspari V, Consolandi C, et al. Diversity of vaginal microbiome and metabolome during genital infections. Sci Rep. 2019;9(1):14095.

37. Chiu SF, Huang PJ, Cheng WH, Huang CY, Chu LJ, Lee CC, et al. Vaginal Microbiota of the Sexually Transmitted Infections Caused by Chlamydia trachomatis and Trichomonas vaginalis in Women with Vaginitis in Taiwan. Microorganisms. 2021;9(9).

38. Cheong HC, Yap PSX, Chong CW, Cheok YY, Lee CYQ, Tan GMY, et al. Diversity of endocervical microbiota associated with genital Chlamydia trachomatis infection and infertility among women visiting obstetrics and gynecology clinics in Malaysia. PLoS One. 2019;14(11):e0224658.

39. Di Pietro M, Filardo S, Porpora MG, Recine N, Latino MA, Sessa R. HPV/Chlamydia trachomatis co-infection: metagenomic analysis of cervical microbiota in asymptomatic women. New Microbiologica. 2018;41(1):34–41.

40. Borgogna JC, Shardell MD, Yeoman CJ, Ghanem KG, Kadriu H, Ulanov AV, et al. The association of Chlamydia trachomatis and Mycoplasma genitalium infection with the vaginal metabolome. Sci Rep. 2020;10(1):3420.

41. Cao K-AL, Boitard S, Besse P. Sparse PLS discriminant analysis: biologically relevant feature selection and graphical displays for multiclass problems. BMC Bioinformatics. 2011;12:1–17.

42. Mallick H, Rahnavard A, McIver LJ, Ma S, Zhang Y, Nguyen LH, et al. Multivariable association discovery in population-scale meta-omics studies. PLoS Comput Biol. 2021;17(11):e1009442.

43. Wang Q, Garrity GM, Tiedje JM, Cole JR. Naive Bayesian classifier for rapid assignment of rRNA sequences into the new bacterial taxonomy. Appl Environ Microbiol. 2007;73(16):5261–7.

44. Poston TB, Lee DE, Darville T, Zhong W, Dong L, O’Connell CM, et al. Cervical Cytokines Associated With Chlamydia trachomatis Susceptibility and Protection. J Infect Dis. 2019;220(2):330–9.

45. González, Déjean, Martin, Baccini. CCA: An R package to extend canonical correlation analysis. Journal of Statistical Software. 2008;23(12):1–14.

46. Tamarelle J, Thiébaut AC, De Barbeyrac B, Bebear C, Ravel J, Delarocque-Astagneau E. The vaginal microbiota and its association with human papillomavirus, Chlamydia trachomatis, Neisseria gonorrhoeae and Mycoplasma genitalium infections: a systematic review and meta-analysis. Clinical Microbiology and Infection. 2019;25(1):35–47.

47. Filardo S, Di Pietro M, Tranquilli G, Latino MA, Recine N, Porpora MG, et al. Selected Immunological Mediators and Cervical Microbial Signatures in Women with Chlamydia trachomatis Infection. mSystems. 2019;4(4).

48. Sasaki-Imamura T, Yoshida Y, Suwabe K, Yoshimura F, Kato H. Molecular basis of indole production catalyzed by tryptophanase in the genus Prevotella. FEMS Microbiology Letters. 2011;322(1):51–9.

49. Buhl M, Dunlap C, Marschal M. Prevotella brunnea sp. nov., isolated from a wound of a patient. International Journal of Systematic and Evolutionary Microbiology. 2019;69(12):3933–8.

50. Horvath A, Durdevic M, Leber B, di Vora K, Rainer F, Krones E, et al. Changes in the Intestinal Microbiome during a Multispecies Probiotic Intervention in Compensated Cirrhosis. Nutrients. 2020;12(6).

51. Hinderfeld AS, Phukan N, Bär A-K, Roberton AM, Simoes-Barbosa A. Cooperative Interactions between Trichomonas vaginalis and Associated Bacteria Enhance Paracellular Permeability of the Cervicovaginal Epithelium by Dysregulating Tight Junctions. Infection and immunity. 2019;87(5).

52. Goldstein EJ, Tyrrell KL, Citron DM. Lactobacillus species: taxonomic complexity and controversial susceptibilities. Clin Infect Dis. 2015;60 Suppl 2:S98–107.

53. O’Callaghan JL, Willner D, Buttini M, Huygens F, Pelzer ES. Limitations of 16S rRNA Gene Sequencing to Characterize Lactobacillus Species in the Upper Genital Tract. Frontiers in Cell and Developmental Biology. 2021;9:641921.

54. Callahan BJ, McMurdie PJ, Holmes SP. Exact sequence variants should replace operational taxonomic units in marker-gene data analysis. ISME J. 2017;11(12):2639–43.

55. Johnson JS, Spakowicz DJ, Hong BY, Petersen LM, Demkowicz P, Chen L, et al. Evaluation of 16S rRNA gene sequencing for species and strain-level microbiome analysis. Nat Commun. 2019;10(1):5029.

56. O’Flaherty S, Briner Crawley A, Theriot CM, Barrangou R. The Lactobacillus Bile Salt Hydrolase Repertoire Reveals Niche-Specific Adaptation. mSphere. 2018;3(3).

57. Navarro S, Abla H, Delgado B, Colmer-Hamood JA, Ventolini G, Hamood AN. Glycogen availability and pH variation in a medium simulating vaginal fluid influence the growth of vaginal Lactobacillus species and Gardnerella vaginalis. BMC Microbiol. 2023;23(1):186.

58. Hertzberger R, May A, Kramer G, van Vondelen I, Molenaar D, Kort R. Genetic Elements Orchestrating Lactobacillus crispatus Glycogen Metabolism in the Vagina. Int J Mol Sci. 2022;23(10).

59. Putonti C, Shapiro JW, Ene A, Tsibere O, Wolfe AJ. Comparative genomic study of Lactobacillus jensenii and the newly defined Lactobacillus mulieris species identifies species-specific functionality. mSphere. 2020;5(4):00560–20.

60. Hood MI, Skaar EP. Nutritional immunity: transition metals at the pathogen-host interface. Nat Rev Microbiol. 2012;10(8):525–37.

61. Raulston JE. Response of Chlamydia trachomatis serovar E to iron restriction in vitro and evidence for iron-regulated chlamydial proteins. Infection and Immunity. 1997;65(11):4539–47.

62. Pokorzynski ND, Thompson CC, Carabeo RA. Ironing Out the Unconventional Mechanisms of Iron Acquisition and Gene Regulation in Chlamydia. Frontiers in Cellular and Infection Microbiology. 2017;7:394.

63. Pokorzynski ND, Alla MR, Carabeo RA. Host cell amplification of nutritional stress contributes to persistence in Chlamydia trachomatis. mBio. 2022;13(6):e02719–22.

64. Pokorzynski ND, Brinkworth AJ, Carabeo R. A bipartite iron-dependent transcriptional regulation of the tryptophan salvage pathway in Chlamydia trachomatis. Elife. 2019;8.

65. Pickering JL, Prosser A, Corscadden KJ, De Gier C, Richmond PC, Zhang G, et al. Haemophilus haemolyticus Interaction with Host Cells Is Different to Nontypeable Haemophilus influenzae and Prevents NTHi Association with Epithelial Cells. Frontiers in cellular and infection microbiology. 2016;6:50.

66. Fulte S, Atto B, McCarty A, Horn KJ, Redzic JS, Eisenmesser E, et al. Heme sequestration by hemophilin from Haemophilus haemolyticus reduces respiratory tract colonization and infection with non-typeable Haemophilus influenzae. mSphere. 2024;9(3):e00006–24.

67. Latham RD, Torrado M, Atto B, Walshe JL, Wilson R, Guss JM, et al. A heme-binding protein produced by Haemophilus haemolyticus inhibits non-typeable Haemophilus influenzae. Mol Microbiol. 2020;113(2):381–98.

68. Woodall CA, Hammond A, Cleary D, Preston A, Muir P, Pascoe B, et al. Oral and gut microbial biomarkers of susceptibility to respiratory tract infection in adults: A feasibility study. Heliyon. 2023;9(8):e18610.

69. Hiippala K, Kainulainen V, Kalliomaki M, Arkkila P, Satokari R. Mucosal Prevalence and Interactions with the Epithelium Indicate Commensalism of Sutterella spp. Front Microbiol. 2016;7:1706.

70. Ceccarani C, Marangoni A, Severgnini M, Camboni T, Laghi L, Gaspari V, et al. Rectal Microbiota Associated With Chlamydia trachomatis and Neisseria gonorrhoeae Infections in Men Having Sex With Other Men. Front Cell Infect Microbiol. 2019;9:358.

71. Atarashi K, Tanoue T, Ando M, Kamada N, Nagano Y, Narushima S, et al. Th17 Cell Induction by Adhesion of Microbes to Intestinal Epithelial Cells. Cell. 2015;163(2):367–80.

72. Gaboriau-Routhiau V, Rakotobe S, Lecuyer E, Mulder I, Lan A, Bridonneau C, et al. The key role of segmented filamentous bacteria in the coordinated maturation of gut helper T cell responses. Immunity. 2009;31(4):677–89.

73. Yount KS, Kollipara A, Liu C, Zheng X, O’Connell CM, Bagwell B, et al. Unique T cell signatures are associated with reduced Chlamydia trachomatis reinfection in a highly exposed cohort. bioRxiv. 2023.

74. Fettweis JM, Serrano MG, Huang B, Brooks JP, Glascock AL, Sheth NU, et al. An emerging mycoplasma associated with trichomoniasis, vaginal infection and disease. PLoS One. 2014;9(10):e110943.

75. Ioannidis A, Papaioannou P, Magiorkinis E, Magana M, Ioannidou V, Tzanetou K, et al. Detecting the Diversity of Mycoplasma and Ureaplasma Endosymbionts Hosted by Trichomonas vaginalis Isolates. Front Microbiol. 2017;8:1188.

76. Ma C, Du J, Dou Y, Chen R, Li Y, Zhao L, et al. The Associations of Genital Mycoplasmas with Female Infertility and Adverse Pregnancy Outcomes: a Systematic Review and Meta-analysis. Reprod Sci. 2021;28(11):3013–31.

77. Jonduo ME, Vallely LM, Wand H, Sweeney EL, Egli-Gany D, Kaldor J, et al. Adverse pregnancy and birth outcomes associated with Mycoplasma hominis, Ureaplasma urealyticum and Ureaplasma parvum: a systematic review and meta-analysis. BMJ Open. 2022;12(8):e062990.

78. Margarita V, Rappelli P, Dessi D, Pintus G, Hirt RP, Fiori PL. Symbiotic association with Mycoplasma hominis can influence growth rate, ATP production, cytolysis and inflammatory response of Trichomonas vaginalis. Frontiers in Microbiology. 2016;7:953.

79. Margarita V, Bailey NP, Rappelli P, Diaz N, Dessì D, Fettweis JM, et al. Two Different Species of Mycoplasma Endosymbionts Can Influence Trichomonas vaginalis Pathophysiology. mBio. 2022;13(3):e00918–22.

80. Fiori PL, Diaz N, Cocco AR, Rappelli P, Dessi D. Association of Trichomonas vaginalis with its symbiont Mycoplasma hominis synergistically upregulates the in vitro proinflammatory response of human monocytes. Sex Transm Infect. 2013;89(6):449–54.

81. Greub G, Raoult D. “Actinobaculum massiliae,” a new species causing chronic urinary tract infection. J Clin Microbiol. 2002;40(11):3938–41.

82. Carrillo-Avila JA, Bonilla-Garcia L, Navarro-Mari JM, Gutierrez-Fernandez J. The first reported case of pelvic inflammatory disease caused by Actinobaculum massiliense. Anaerobe. 2019;55:93–5.

83. Yu Y, Tsitrin T, Singh H, Doerfert SN, Sizova MV, Epstein SS, et al. Actinobaculum massiliense Proteome Profiled in Polymicrobial Urethral Catheter Biofilms. Proteomes. 2018;6(4).

84. Reynoso-García J, Miranda-Santiago AE, Meléndez-Vázquez NM, Acosta-Pagán K, Sánchez-Rosado M, Díaz-Rivera J, et al. A complete guide to human microbiomes: Body niches, transmission, development, dysbiosis, and restoration. Frontiers in Systems Biology. 2022;2:951403.

85. Lamas A, Regal P, Vázquez B, Cepeda A, Franco CM. Short chain fatty acids commonly produced by gut microbiota influence Salmonella enterica motility, biofilm formation, and gene expression. Antibiotics. 2019;8(4):265.

86. Ozma MA, Abbasi A, Akrami S, Lahouty M, Shahbazi N, Ganbarov K, et al. Postbiotics as the key mediators of the gut microbiota-host interactions. Infez Med. 2022;30(2):180–93.

87. Mirzaei R, Kavyani B, Nabizadeh E, Kadkhoda H, Asghari Ozma M, Abdi M. Microbiota metabolites in the female reproductive system: Focused on the short-chain fatty acids. Heliyon. 2023;9(3):e14562.

88. Wiesenfeld HC, Sweet RL, Ness RB, Krohn MA, Amortegui AJ, Hillier SL. Comparison of Acute and Subclinical Pelvic Inflammatory Disease. Sexually transmitted diseases. 2005;32(7):400–5.

89. Wiesenfeld HC, Hillier SL, Meyn LA, Amortegui AJ, Sweet RL. Subclinical pelvic inflammatory disease and infertility. Obstet Gynecol. 2012;120(1):37–43.

90. Dodet B. Current barriers, challenges and opportunities for the development of effective STI vaccines: point of view of vaccine producers, biotech companies and funding agencies. Vaccine. 2014;32(14):1624–9.

91. Broutet N, Fruth U, Deal C, Gottlieb SL, Rees H. Vaccines against sexually transmitted infections: the way forward. Vaccine. 2014;32(14):1630–7.

92. Poston TB, Gottlieb SL, Darville T. Status of vaccine research and development of vaccines for Chlamydia trachomatis infection. Vaccine. 2019;37(50):7289–94.

93. Workowski KA, Bolan GA, Prevention CfDCa. Sexually transmitted diseases treatment guidelines, 2015. MMWR Recomm Rep. 2015;64(3).

94. Zheng X, O’Connell CM, Zhong W, Poston TB, Wiesenfeld HC, Hillier SL, et al. Gene Expression Signatures Can Aid Diagnosis of Sexually Transmitted Infection-Induced Endometritis in Women. Front Cell Infect Microbiol. 2018;8:307.

95. O’Connell CM, Brochu H, Girardi J, Harrell E, Jones A, Darville T, et al. Simultaneous profiling of sexually transmitted bacterial pathogens, microbiome, and concordant host response in cervical samples using whole transcriptome sequencing analysis. Microb Cell. 2019;6(3):177–83.

96. Abraham S, Juel HB, Bang P, Cheeseman HM, Dohn RB, Cole T, et al. Safety and immunogenicity of the chlamydia vaccine candidate CTH522 adjuvanted with CAF01 liposomes or aluminium hydroxide: a first-in-human, randomised, double-blind, placebo-controlled, phase 1 trial. The Lancet infectious diseases. 2019;19(10):1091–100.

97. Pan P, Gu Y, Sun D-L, Wu QL, Zhou N-Y. Microbial diversity biased estimation caused by intragenomic heterogeneity and interspecific conservation of 16s rrna genes. Applied and Environmental Microbiology. 2023;89(5):15.

98. Kembel SW, Wu M, Eisen JA, Green JL. Incorporating 16S gene copy number information improves estimates of microbial diversity and abundance. PLoS Comput Biol. 2012;8(10):e1002743.

99. Klappenbach JA, Saxman PR, Cole JR, Schmidt TM. rrndb: the Ribosomal RNA Operon Copy Number Database. Nucleic Acids Research. 2001;29(1):181–4.

100. Douglas GM, Maffei VJ, Zaneveld JR, Yurgel SN, Brown JR, Taylor CM, et al. PICRUSt2 for prediction of metagenome functions. Nat Biotechnol. 2020;38:685–8.

101. Miao J, Chen T, Misir M, Lin Y. Deep learning for predicting 16S rRNA gene copy number. Sci Rep. 2024;14(1):14282.

102. Angly FE, Dennis PG, Skarshewski A, Vanwonterghem I, Hugenholtz P, Tyson GW. CopyRighter: a rapid tool for improving the accuracy of microbial community profiles through lineage-specific gene copy number correction. Microbiome. 2014;2(11).

103. Stoddard SF, Smith BJ, Hein R, Roller BR, Schmidt TM. rrnDB: improved tools for interpreting rRNA gene abundance in bacteria and archaea and a new foundation for future development. Nucleic Acids Res. 2015;43(Database issue):D593–8.

104. Liu C, Hufnagel K, O’Connell CM, Goonetilleke N, Mokashi N, Waterboer T, et al. Reduced Endometrial Ascension and Enhanced Reinfection Associated With Immunoglobulin G Antibodies to Specific Chlamydia trachomatis Proteins in Women at Risk for Chlamydia. J Infect Dis. 2022;225(5):846–55.

105. Apprill A, McNally S, Parsons R, Weber L. Minor revision to V4 region SSU rRNA 806R gene primer greatly increases detection of SAR11 bacterioplankton. Aquat Microb Ecol. 2015;75:129–37.

106. Parada AE, Needham DM, Fuhrman JA. Every base matters: assessing small subunit rRNA primers for marine microbiomes with mock communities, time series and global field samples. Environ Microbiol. 2016;18(5):1403–14.

107. Edgar RC. UNOISE2: improved error-correction for Illumina 16S and ITS amplicon sequencing. BioRxiv. 2016:081257.

108. Zhang J, Kobert K, Flouri T, Stamatakis A. PEAR: a fast and accurate Illumina Paired-End reAd mergeR. Bioinformatics. 2014;30(5):614–20.

109. Antich A, Palacin C, Wangensteen OS, Turon X. To denoise or to cluster, that is not the question: optimizing pipelines for COI metabarcoding and metaphylogeography. BMC Bioinformatics. 2021;22(1):177.

110. Li H. Aligning sequence reads, clone sequences and assembly contigs with BWA-MEM. arXiv. 2013.

111. Edgar RC. MUSCLE: multiple sequence alignment with high accuracy and high throughput. Nucleic acids research. 2004;32(5):1792–7.

112. Okonechnikov K, Golosova O, Fursov M. Unipro UGENE: a unified bioinformatics toolkit. Bioinformatics. 2012;28(8):1166–7.

113. Edgar RC. SINTAX: a simple non-Bayesian taxonomy classifier for 16S and ITS sequences. biorxiv. 2016.

114. Antonio MAD, Hawes SE, Hillier SL. The Identification of Vaginal Lactobacillus Species and the Demographic and Microbiologic Characteristics of Women Colonized by These Species. Journal of Infectious Diseases. 1999;180(6):The Journal of Infectious Diseases.

115. Gloor GB, Macklaim JM, Pawlowsky-Glahn V, Egozcue JJ. Microbiome Datasets Are Compositional: And This Is Not Optional. Front Microbiol. 2017;8:2224.

116. Aitchison J. The statistical analysis of compositional data. Royal Statistical Society. 1982;44(2):139–77.

117. Quinn TP, Erb I, Gloor G, Notredame C, Richardson MF, Crowley TM. A field guide for the compositional analysis of any-omics data. Gigascience. 2019;8(9).

118. France MT, Ma B, Gajer P, Brown S, Humphrys MS, Holm JB, et al. VALENCIA: a nearest centroid classification method for vaginal microbial communities based on composition. Microbiome. 2020;8(1):166.

119. Rohart F, Gautier B, Singh A, Le Cao KA. mixOmics: An R package for ‘omics feature selection and multiple data integration. PLoS Comput Biol. 2017;13(11):e1005752.

120. Liaw A, Wiener M. Classification and Regression by randomForest. R News. 2002;2(3):18–22.

121. Robin X, Turck N, Hainard A, Tiberti N, Lisacek F, Sanchez J-C, et al. pROC: an open-source package for R and S+ to analyze and compare ROC curves. BMC Bioinformatics. 2011;12:77.

