## Supplementary Materials for "Cervicovaginal microbial features predict *Chlamydia trachomatis* spread to the upper genital tract of infected women"

Jeong, et al.

| Page # | Supplementary Figures / Table |
| --- | --- |
| 2 | Supplementary Figure 1. Overview of cervical microbiome profiles obtained from TRAC cohort. |
| 3 | Supplementary Figure 2. Discrimination of chlamydial infection status at cervix or endometrium stratified by CST types using sPLS-DA components. |
| 4 | Supplementary Figure 3. Distribution of *Lactobacillus* sp. variants across cohort with CST types. |
| 5 | Supplementary Figure 4. Multiple alignment of low-abundant *Lactobacillus* ASV sequences differentially abundant between CT+ and CT- groups. |
| 6 | Supplementary Figure 5. ROC curves and corresponding AUCs from random forest prediction for absent CT infection. |
| 7 | Supplementary Figure 6. Distribution of prediction probabilities for four false negatives in a proof-of-concept analysis. |
| 8 | Supplementary Figure 7. Data analysis flowchart. |
| 9 | Supplementary Figure 8. Random Forest classification with K-fold CV. |
| 10 | Supplementary Table S1. The demographic, clinical and biological information of TRAC cohort. |
| 11 | Supplementary Table S2. The associations of CST assignment with CT infection outcomes and race of TRAC cohort. |
| 12 | Supplementary Table S3. The ASVs stably selected for the discrimination between CT+ and CT- women across validation folds across repeats by sPLS-DA. |
| 13 | Supplementary Table S4. Significant ASVs having differential abundances between CT+ and CT- women^[1]^. |
| 14 | Supplementary Table S5. ASVs are predictive of absent CT infection. |
| 15 | Supplementary Table S6. The ASVs consistently chosen for the discrimination between Endo+ and Endo- women by sPLS-DA^[2]^. |
| 16 | Supplementary Table S7. Overview of RDP Classifier 16S trainset No18 raw training database. |

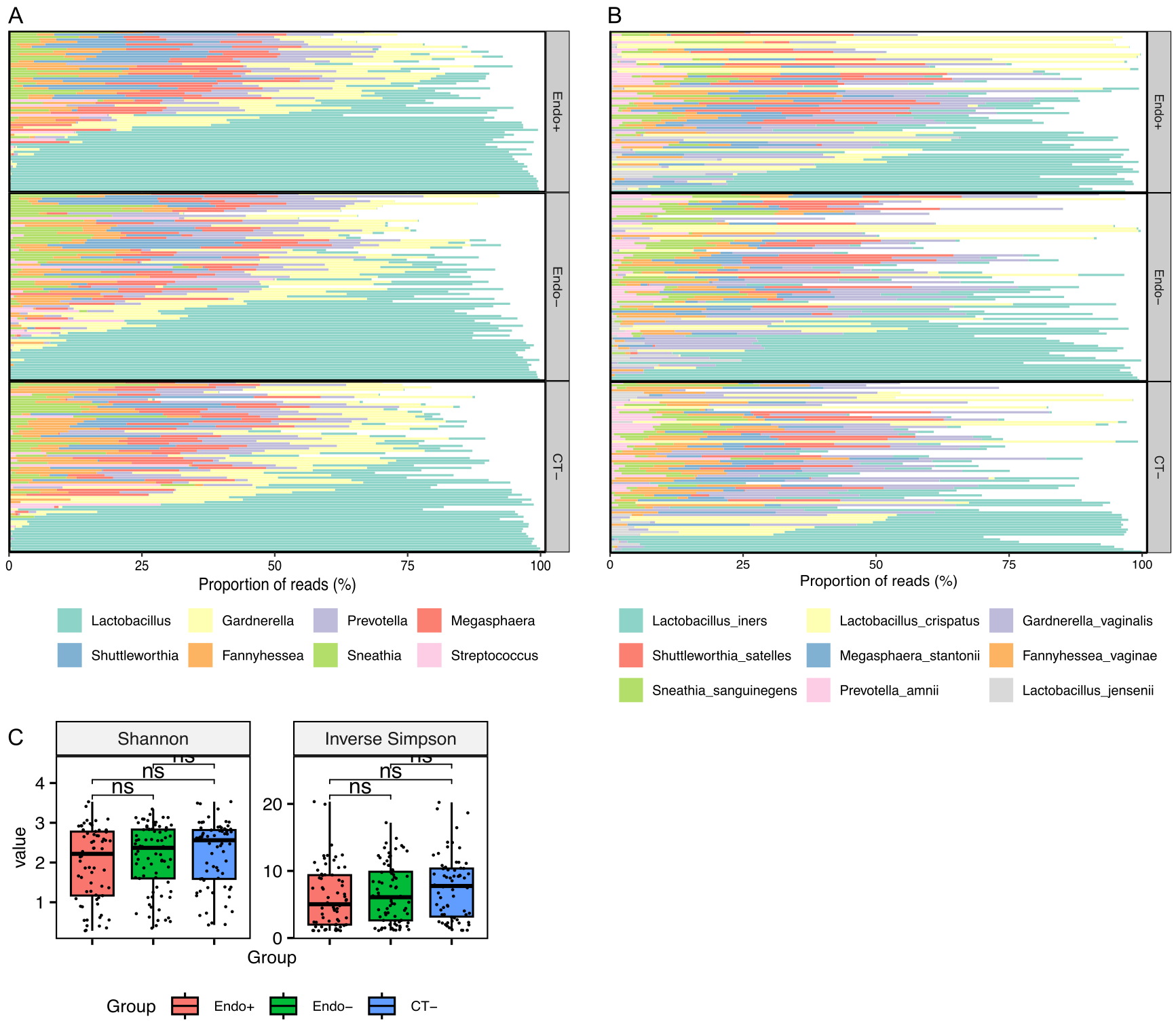

**Supplementary Figure 1. Overview of cervical microbiome profiles obtained from TRAC cohort. A.** The relative abundances of genus in each woman are represented in stacked bars for three groups (Endo+, Endo-, and CT-). The genus labels are arranged based on the most abundant genus across samples. **B.** The relative abundances of species in each woman are shown as stacked bars for three groups. The species labels were ordered in the most abundant species across samples. **C.** The alpha diversity indices (Shannon and Inverse Simpson) were derived from cervicovaginal microbiomes across three groups. The pairwise comparisons in the diversity indices between groups were statistically determined using Wilcoxon rank sum tests, and Inverse Simpson diversity between Endo+ and CT- groups was weakly different (* P < 0.05; ns = not significant).

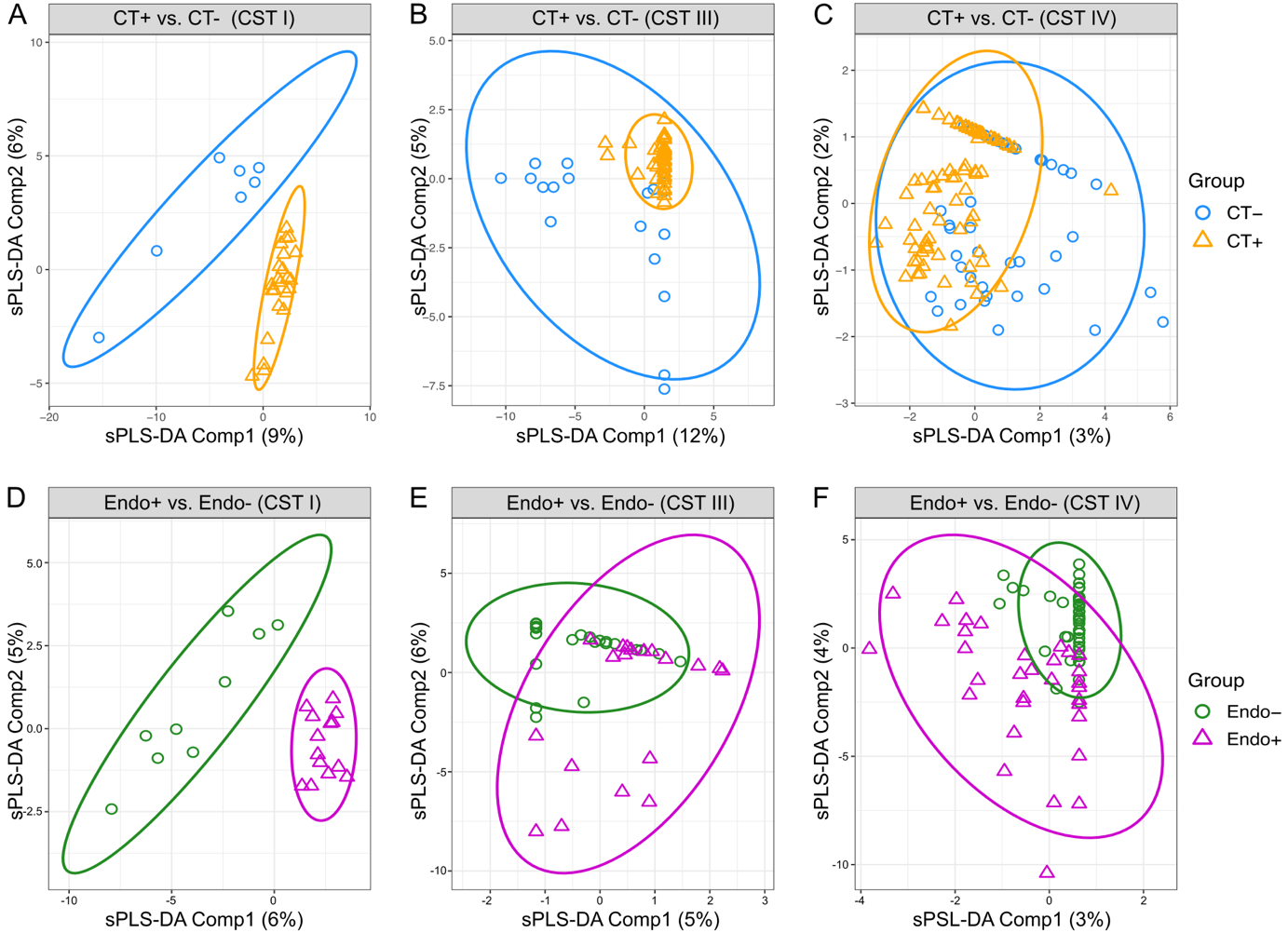

**Supplementary Figure 2. Discrimination of chlamydial infection status at cervix or endometrium stratified by CST types using sPLS-DA components.** Chlamydial infection statuses at cervix or endometrium were discriminated within CST I, CST III, and CST IV, using the first two sPLS-DA components. Discrimination of CT+ from CT- women were performed based on ASVs excluding CT ASV (A, B, and C), while discriminating Endo+ from Endo- women was performed based on ASVs including CT ASV (D, E, and F). The performances of discrimination were evaluated by the mean of AUC. **A.** The first two sPLS-DA components discriminate CT+ (n = 23) from CT- (n = 7) women within CST I (AUC mean = 0.49, AUC s.d. = 0.09, n = 30). **B.** The first two sPLS-DA components discriminate CT+ (n = 43) from CT- (n = 20) women within CST III (AUC mean = 0.71, AUC s.d. = 0.04, n = 63). **C.** The first two sPLS-DA components discriminate CT+ (n = 76) from CT- (n = 41) women within CST IV (AUC mean = 0.50, AUC s.d. = 0.04, n = 117). **D.** The first two sPLS-DA components discriminate Endo+ (n = 14) from Endo- (n = 9) women within CST I (AUC mean = 0.47, AUC s.d. = 0.10, n = 23). **E.** The first two sPLS-DA components discriminate Endo+ (n = 19) from Endo- (n = 24) women within CST III (AUC mean = 0.58, AUC s.d. = 0.09, n = 43). **F.** The first two sPLS-DA components discriminate Endo+ (n = 32) from Endo- (n = 44) women within CST IV (AUC mean = 0.71, AUC s.d. = 0.03, n = 76).

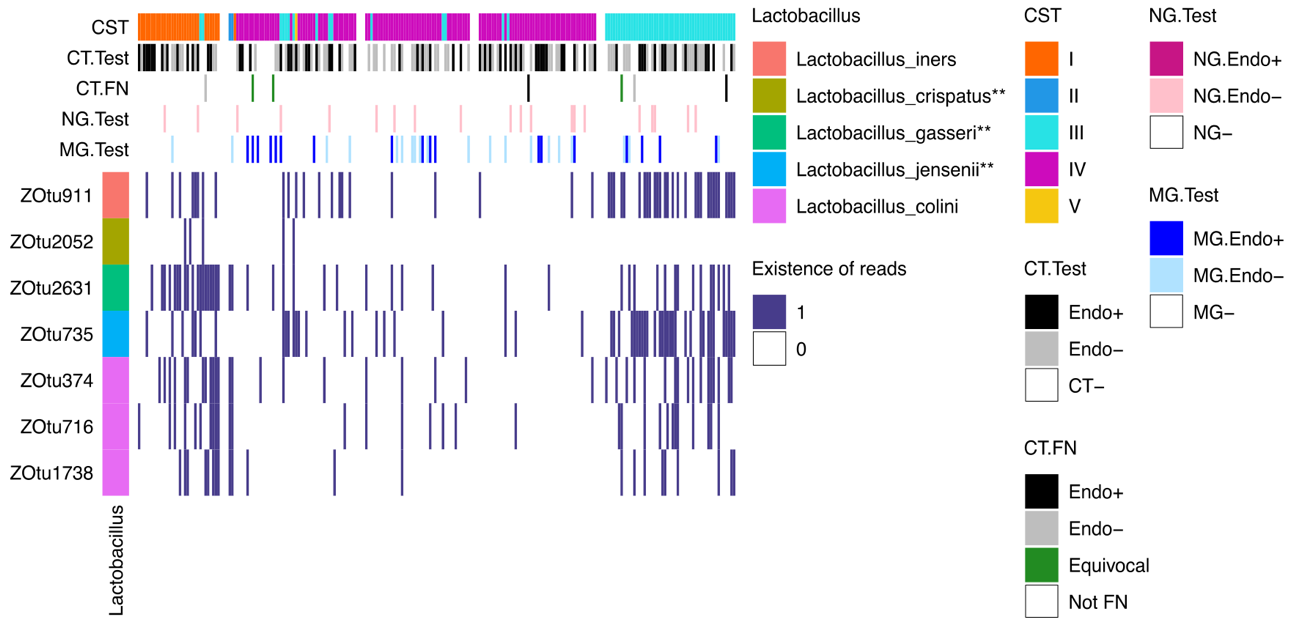

**Supplementary Figure 3. Distribution of *Lactobacillus* sp. variants across cohort with CST types.** The heatmap illustrates the presence of 7 ASVs of *Lactobacillus sp.* that were identified as significantly differentially abundant between CT+ and CT- women (**Supplementary Table S3**; unadjusted p-value < 0.05). The *Lactobacillus* variants, identified through the 7 ASV sequences, were low-abundant but significantly associated with CT infection at cervix. The samples were clustered in the same order of heatmap in **Figure 1A**.

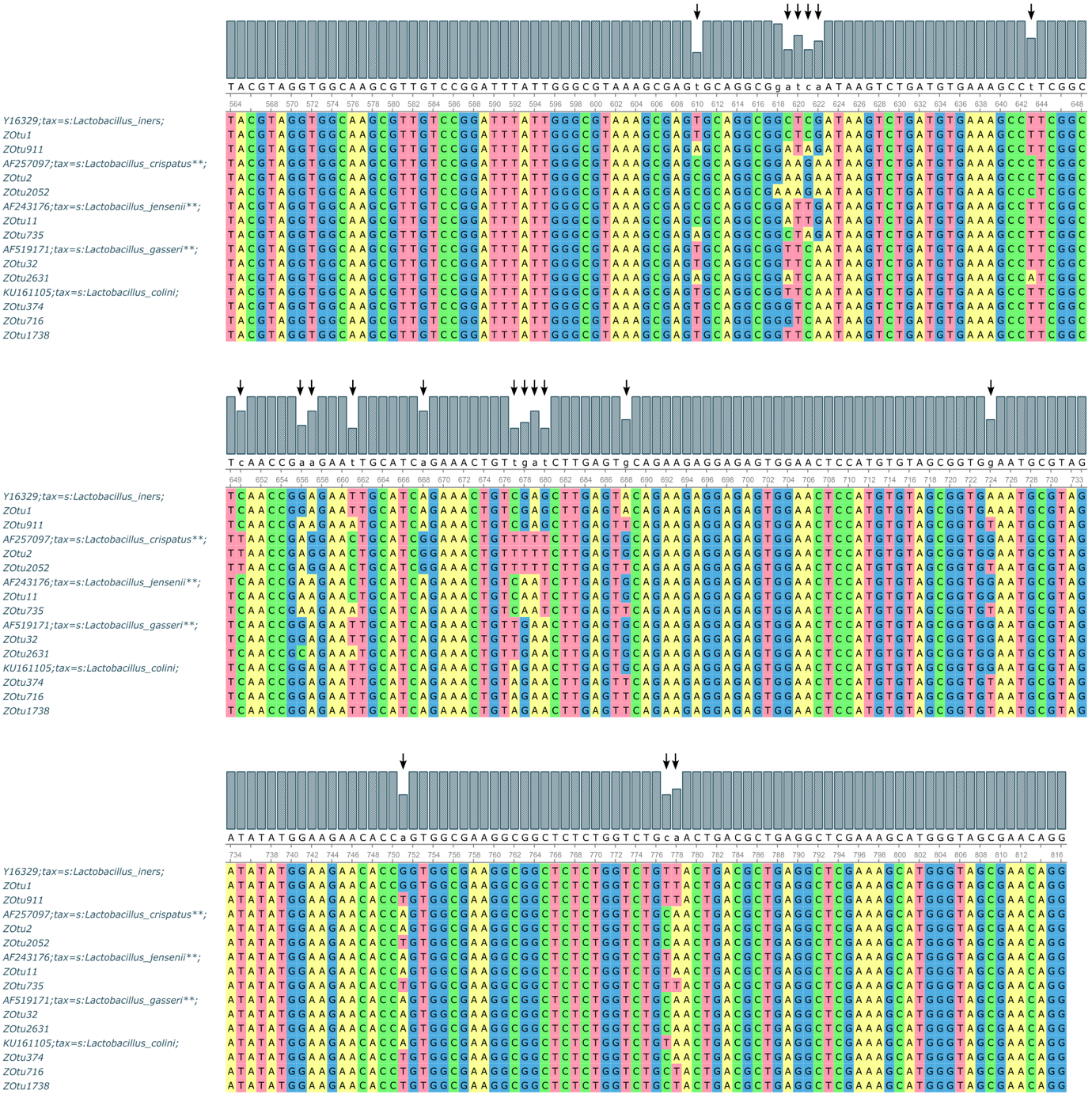

**Supplementary Figure 4. Multiple alignment of low-abundant *Lactobacillus* ASV sequences differentially abundant between CT+ and CT- groups.** The 7 *Lactobacillus* ASVs, identified as low-abundant but significantly differentially abundant between CT+ and CT- groups, are listed in **Supplementary Table S3**. These Lactobacillus ASVs were aligned with the corresponding ASVs of their most abundant Lactobacillus ASV sequences and their reference sequences of 16S V4 region obtained from RDP 16S No18 reference database. Arrows above the alignments indicate 20 positions of frequent variants. The alignment suggests that these low-abundant *Lactobacillus* ASV sequences were unlikely generated by sequencing errors.

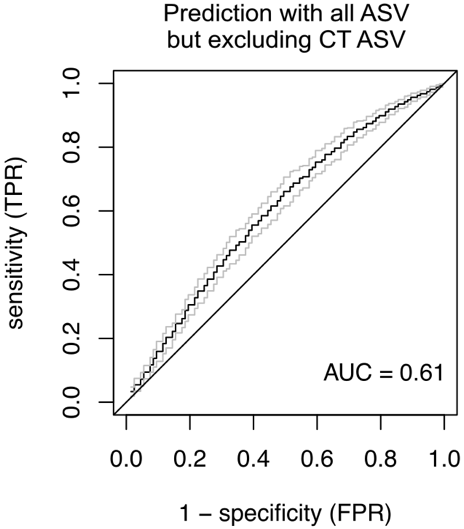

**Supplementary Figure 5. ROC curves and corresponding AUCs from random forest prediction for absent CT infection.** Prediction accuracies were evaluated by averaging the area under the curves (AUCs) over 100 replicates using ASVs without CT ASV. The prediction performance based on ASVs excluding CT ASV, was illustrated by averaged receiver operating characteristic (ROC) curves, having an AUC of 0.61.

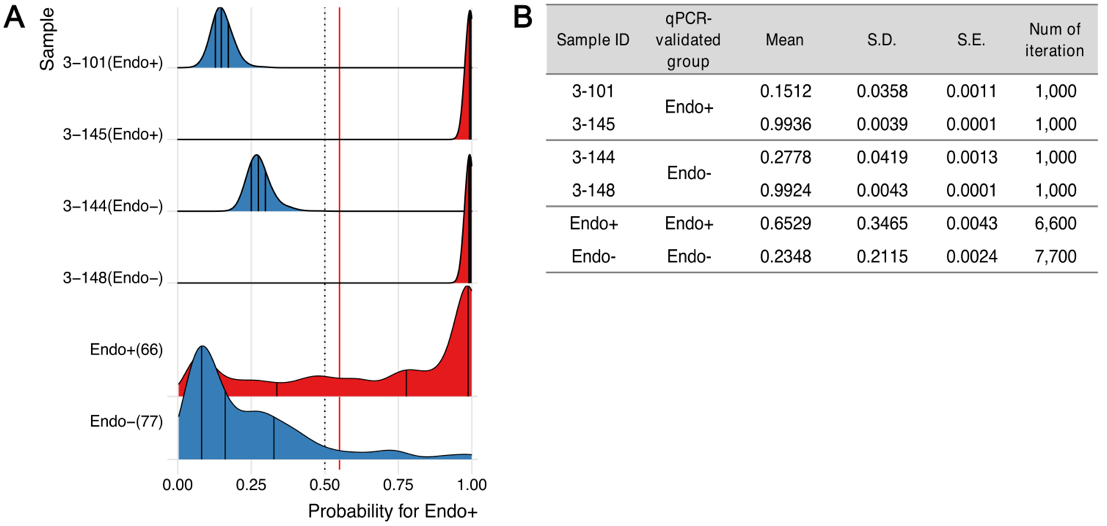

**Supplementary Figure 6. Distribution of prediction probabilities for four false negatives in a proof-of-concept analysis.** The prediction probabilities of chlamydial ascension range from 0 to 1, with higher values indicating a higher risk of chlamydial ascension. In A and C figures, the samples predicted to be Endo+ were colored red, while those predicted to be Endo- were colored blue. **A.** The prediction probabilities for seven women were assessed using random-forest prediction based on 12 informative ASVs. The red vertical line indicates the optimal threshold of 0.55, determined by prediction analysis. **B.** The mean, standard deviation (S.D.), and standard error (S.E.) of the Endo+ probabilities represent the average, variability, and precision, respectively.

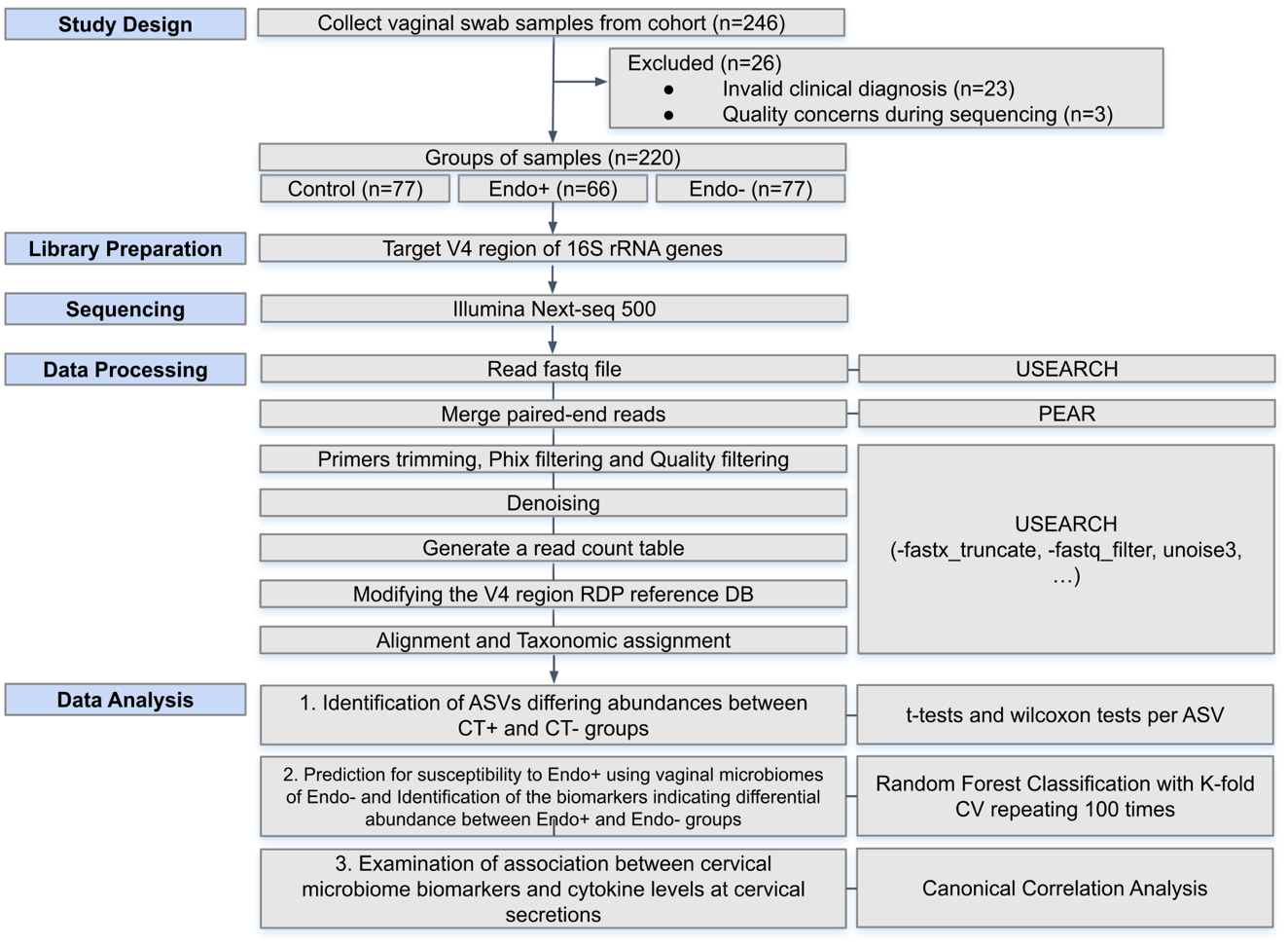

**Supplementary Figure 7. Data analysis flowchart.** The pipeline includes the entire process from sample collection and 16S sequencing to data analysis. For each step of computational data processing, the corresponding bioinformatics tools are indicated on the right-hand side. The primary statistical analysis methods are specified for the three main steps of data analysis.

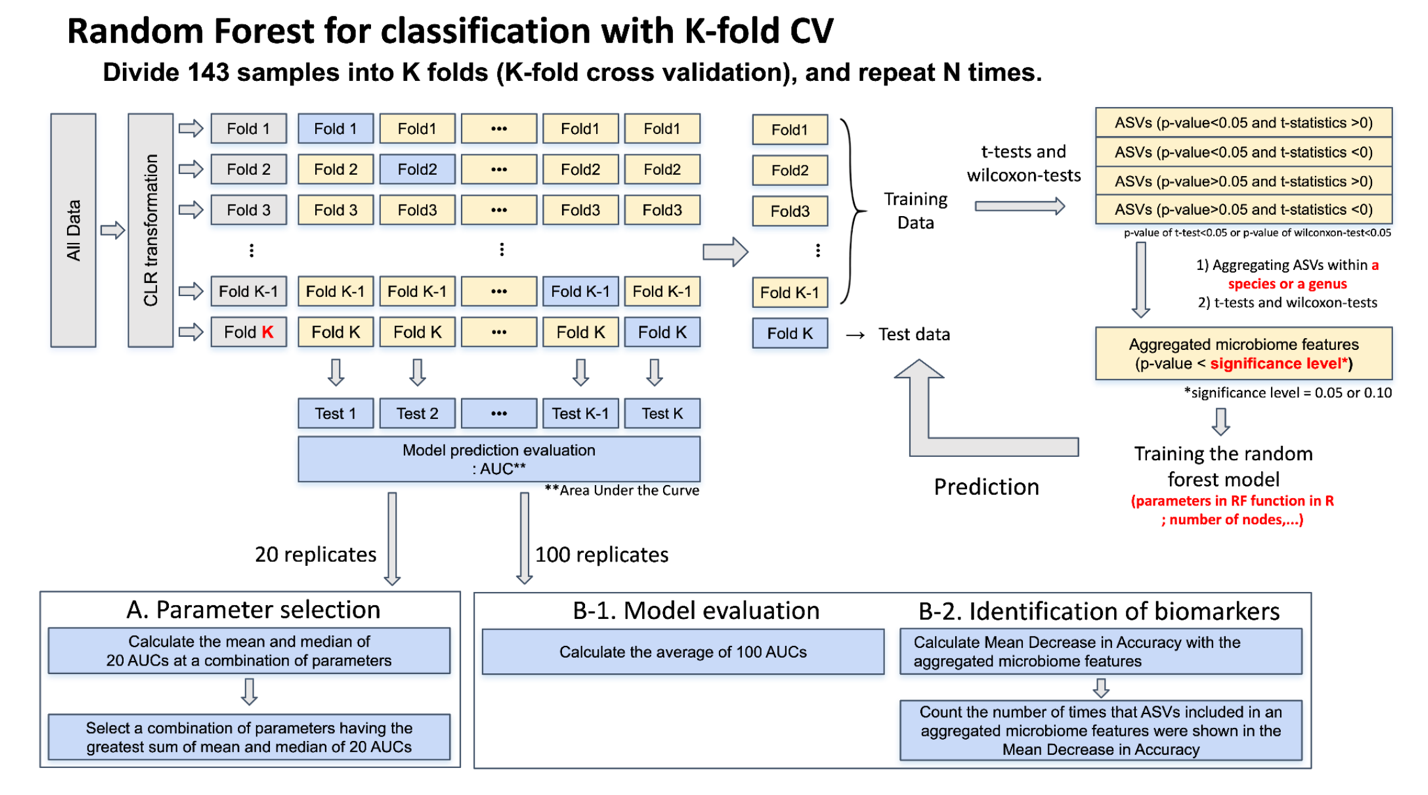

**Supplementary Figure 8. Random Forest classification with K-fold CV. A.** The dataset was split into training set and test set using K-fold CV and then replicated with 20 times for parameter selection. The red highlighted parameters were determined after A session. **B-1.** With the selected parameters, the prediction model was assessed averaging AUCs with 100 times for robust results of random forest prediction. **B-2.** The most informative ASVs contributing to the prediction were selected using “mean decrease in accuracy” as the metric for feature importance.

**Supplementary Table S1. The demographic, clinical and biological information of TRAC cohort.** For data analysis, 220 out of 246 samples from the TRAC cohort were utilized, and their demographic, clinical, and biological information is summarized. Among 220 women, seven false negatives were excluded from CT- group, and separately summarized. The remaining 213 women were classified into three groups (CT-, Endo-, and Endo+), with their characteristics displayed for each group in the respective rows. Classification into CT infection groups and co-infection results were based on clinical diagnostic tests, while CT ASV detection and CST were based on 16S data. The other information was obtained through a questionnaire at enrollment.

|  |  |  | n (% in group) | | |  | n | | |
| --- | --- | --- | --- | --- | --- | --- | --- | --- | --- |
|  |  |  | Clinical Testing results (n=213) | | |  | qPCR-validated group (n=7) | | |
| Attributes |  | Sum (n) | Endo+ (n=66) | Endo- (n=77) | CT- (n=70) |  | Endo+ (n=2) | Endo- (n=2) | Equivocal (n=3) |
| Age | 18-20 | 86 | 36 (54.5) | 38 (49.4) | 12 (17.1) |  | 1 | 1 | 3 |
|  | 21-25 | 97 | 24 (36.4) | 31 (40.3) | 42 (60.0) |  | 1 | 1 |  |
|  | 26-30 | 21 | 5 (7.6) | 5 (6.5) | 11 (15.7) |  |  |  |  |
|  | 31-35 | 8 | 0 (0) | 3 (3.9) | 5 (7.1) |  |  |  |  |
|  | N/A | 1 | 1 (1.5) | 0 (0) | 0 (0) |  |  |  |  |
| Race | Asian | 1 | 1 (1.5) | 0 (0) | 0 (0) |  |  |  |  |
|  | Bi/Multi-racial | 29 | 8 (12.1) | 12 (15.6) | 9 (12.9) |  |  |  | 1 |
|  | Black/AA | 134 | 41 (62.1) | 51 (66.2) | 42 (60.0) |  | 2 | 1 | 2 |
|  | Caucasian/White | 44 | 14 (21.2) | 13 (16.9) | 17 (24.3) |  |  | 1 |  |
|  | Other | 5 | 2 (3.0) | 1 (1.3) | 2 (2.9) |  |  |  |  |
| CST types | I | 30 | 14 (21.2) | 9 (11.7) | 7 (10.0) |  |  | 1 |  |
|  | II | 2 | 0 (0) | 0 (0) | 2 (2.9) |  |  |  |  |
|  | III | 63 | 19 (28.8) | 24 (31.2) | 20 (28.6) |  | 1 | 1 | 1 |
|  | IV | 117 | 32 (48.5) | 44 (57.1) | 41 (58.6) |  | 1 |  | 2 |
|  | V | 1 | 1 (1.5) | 0 (0) | 0 (0) |  |  |  |  |
| CT ASV detection | Yes | 83 | 53 (80.3) | 30 (39.0) | 0 (0) |  | 2 | 2 | 3 |
|  | No | 130 | 13 (19.7) | 47 (61.0) | 70 (100) |  |  |  |  |
| Co-infection | NG.Endo+ | 0 | 0 (0) | 0 (0) | 0 (0) |  |  |  |  |
|  | NG.Endo- | 20 | 11 (16.7) | 8 (10.4) | 1 (1.4) |  |  |  |  |
|  | NG- | 193 | 55 (83.3) | 69 (89.6) | 69 (98.6) |  | 2 | 2 | 3 |
|  | MG.Endo+ | 17 | 7 (10.6) | 5 (6.5) | 5 (7.1) |  |  |  | 1 |
|  | MG.Endo- | 20 | 2 (3.0) | 14 (18.2) | 4 (5.7) |  |  |  |  |
|  | MG- | 176 | 57 (86.4) | 58 (75.3) | 61 (87.1) |  | 2 | 2 | 2 |
| Nugent score | Normal (0-3) | 57 | 19 (28.8) | 21 (27.3) | 17 (24.3) |  | 1 | 1 |  |
|  | Intermediate (4-6) | 41 | 12 (18.2) | 15 (19.5) | 14 (20.0) |  |  |  | 1 |
|  | BV (7-10) | 114 | 35 (53.0) | 41 (53.2) | 38 (54.3) |  | 1 | 1 | 2 |
|  | N/A | 1 | 0 (0.0) | 0 (0.0) | 1 (1.4) |  |  |  |  |
| Endometritis by histology | Acute | 7 | 2 (3.0) | 4 (5.2) | 1 (1.4) |  |  |  |  |
|  | Chronic | 35 | 14 (21.2) | 10 (13.0) | 11 (15.7) |  | 1 | 1 | 2 |
|  | Insufficient tissue | 73 | 18 (27.3) | 34 (44.2) | 21 (30.0) |  |  |  |  |
|  | No | 80 | 26 (39.4) | 23 (29.9) | 31 (44.3) |  | 1 | 1 | 1 |
|  | N/A | 18 | 6 (9.1) | 6 (7.8) | 6 (8.6) |  |  |  |  |
| IUD/Depo | IUD | 24 | 8 (12.1) | 9 (11.7) | 7 (10.0) |  |  | 1 | 1 |
|  | Depo | 25 | 8 (12.1) | 13 (16.9) | 4 (5.7) |  |  |  | 1 |
|  | No | 163 | 49 (74.2) | 55 (71.4) | 59 (84.3) |  | 2 | 1 | 1 |
|  | N/A | 1 | 1 (1.5) | 0 (0) | 0 (0) |  |  |  |  |
| Days since last menstrual period | <= 14 days | 82 | 24 (36.4) | 26 (33.8) | 32 (45.7) |  | 1 |  |  |
|  | <= 31 days | 76 | 28 (42.4) | 24 (31.2) | 24 (34.3) |  |  |  | 1 |
|  | <= 62 days | 16 | 3 (4.5) | 7 (9.1) | 6 (8.6) |  | 1 |  |  |
|  | > 62 days | 35 | 9 (13.6) | 18 (23.4) | 8 (11.4) |  |  | 2 | 1 |
|  | Unsure | 2 | 0 (0) | 2 (2.6) | 0 (0) |  |  |  |  |
|  | N/A | 2 | 2 (3.0) | 0 (0) | 0 (0) |  |  |  | 1 |
| Sex during period | Yes | 26 | 10 (15.2) | 2 (2.6) | 14 (20.0) |  |  |  |  |
|  | No | 179 | 55 (83.3) | 71 (92.2) | 53 (75.7) |  | 2 | 1 | 3 |
|  | N/A | 8 | 1 (1.5) | 4 (5.2) | 3 (4.3) |  |  | 1 |  |

**Supplementary Table S2. The associations of CST assignment with CT infection outcomes and race of TRAC cohort.** The associations of CSTs with Cervical infection, endometrial infection and race were examined by Fisher’s exact test. Race exhibited a significant association with CSTs (p-value < 0.05).

| n (% in each attribute category) | | | | | | | | |
| --- | --- | --- | --- | --- | --- | --- | --- | --- |
|  |  |  | Community state types | | | | |  |
| Attributes |  | n | I | II | III | IV | V | P-value |
| Cervical infection | CT+ | 143 | 23 (16) | 0 (0) | 43 (30) | 76 (53) | 1 (1) | 0.2094 |
|  | CT- | 70 | 7 (10) | 2 (3) | 20 (29) | 41 (59) | 0 (0) |  |
| Nugent score level | Normal (0-3) | 59 | 26 (44) | 0 (0) | 31 (53) | 1 (2) | 1 (2) | 0.0005 |
|  | Intermediate (4-6) | 42 | 3(7) | 2 (5) | 26 (62) | 11 (26) | 0 (0) |  |
|  | BV (7-10) | 118 | 2 (2) | 0 (0) | 9 (8) | 107 (91) | 0 (0) |  |
| Race | Asian | 1 | 1 (100) | 0 (0) | 0 (0) | 0 (0) | 0 (0) | 0.01899 |
|  | Bi/Multiple | 30 | 4 (13) | 0 (0) | 9 (30) | 17 (57) | 0 (0) |  |
|  | Black/AA | 139 | 13 (9) | 0 (0) | 44 (32) | 81 (58) | 1 (1) |  |
|  | Caucasian/White | 45 | 12 (27) | 1 (2) | 12 (27) | 20 (44) | 0 (0) |  |
|  | Other | 5 | 1 | 1 | 1 | 2 | 0 |  |

**Supplementary Table S3. The ASVs stably selected for the discrimination between CT+ and CT- women across validation folds across repeats by sPLS-DA.** The table lists ASVs that significantly contributed to the first two components of sPLS-DA, discriminating CT+ and CT- groups across all women (“All”) and within CST types (“CST I”, “CST III”, and “CST IV”). Their contribution was assessed based on frequencies, which were calculated as the proportion of cross-validation folds across repeats. ASVs with a frequency greater than 0.80 for each component were listed.

| CST | Comp | Num | ASV | Freq | Genus | Species | Confidence level | |
| --- | --- | --- | --- | --- | --- | --- | --- | --- |
|  |  |  |  |  |  |  | Genus | Species |
| All | Comp1 | 1 | ZOtu304 | 0.9967 | *Prevotella* | *Prevotella_oris* | 1.0000 | 1.0000 |
|  |  | 2 | ZOtu95 | 0.9599 | *Neisseria* | *Neisseria_gonorrhoeae* | 1.0000 | 0.8900 |
| CST_I | Comp1 | 1 | ZOtu1263 | 1.0000 | *Lactobacillus* | *Lactobacillus_crispatus* | 0.9900 | 0.5148 |
|  |  | 2 | ZOtu443 | 0.9870 | *Lactobacillus* | *Lactobacillus_crispatus* | 0.9600 | 0.4224 |
|  |  | 3 | ZOtu964 | 0.9806 | *Megasphaera* | *Megasphaera_stantonii* | 0.5465 | 0.3661 |
|  |  | 4 | ZOtu548 | 0.9702 | *Megasphaera* | *Megasphaera_stantonii* | 0.3121 | 0.1498 |
|  |  | 5 | ZOtu522 | 0.9696 | *Lactobacillus* | *Lactobacillus_delbrueckii* | 0.8900 | 0.1958 |
|  |  | 6 | ZOtu428 | 0.9382 | *Alloscardovia* | *Alloscardovia_omnicolens* | 1.0000 | 0.9700 |
|  |  | 7 | ZOtu773 | 0.9258 | *Lactobacillus* | *Lactobacillus_crispatus* | 0.9700 | 0.4656 |
|  |  | 8 | ZOtu961 | 0.9170 | *Megasphaera* | *Megasphaera_stantonii* | 0.4467 | 0.3171 |
|  |  | 9 | ZOtu1469 | 0.9094 | *Lactobacillus* | *Lactobacillus_jensenii* | 0.7301 | 0.1387 |
|  |  | 10 | ZOtu2052 | 0.8932 | *Lactobacillus* | *Lactobacillus_crispatus* | 0.9900 | 0.5247 |
|  |  | 11 | ZOtu1288 | 0.8848 | *Lactobacillus* | *Lactobacillus_crispatus* | 0.9100 | 0.3367 |
|  |  | 12 | ZOtu2385 | 0.8764 | *Megasphaera* | *Megasphaera_stantonii* | 0.7316 | 0.4975 |
|  |  | 13 | ZOtu1454 | 0.8618 | *Lactobacillus* | *Lactobacillus_iners* | 0.9600 | 0.8544 |
|  |  | 14 | ZOtu1377 | 0.8602 | *Lactobacillus* | *Lactobacillus_crispatus* | 0.9800 | 0.5488 |
|  |  | 15 | ZOtu962 | 0.8520 | *Megasphaera* | *Megasphaera_stantonii* | 0.3198 | 0.1695 |
|  |  | 16 | ZOtu5 | 0.8514 | *Megasphaera* | *Megasphaera_stantonii* | 0.8022 | 0.4974 |
|  | Comp2 | 1 | ZOtu302 | 0.8082 | *Lactobacillus* | *Lactobacillus_crispatus* | 0.9500 | 0.3610 |
| CST_III | Comp1 | 1 | ZOtu702 | 0.9988 | *Lactobacillus* | *Lactobacillus_crispatus* | 0.9300 | 0.3720 |
|  |  | 2 | ZOtu2139 | 0.9982 | *Lactobacillus* | *Lactobacillus_crispatus* | 0.9800 | 0.4312 |
|  |  | 3 | ZOtu2363 | 0.9930 | *Lactobacillus* | *Lactobacillus_crispatus* | 0.9800 | 0.5390 |
|  |  | 4 | ZOtu1170 | 0.9868 | *Lactobacillus* | *Lactobacillus_crispatus* | 0.8700 | 0.2262 |
|  |  | 5 | ZOtu946 | 0.9476 | *Lactobacillus* | *Lactobacillus_crispatus* | 0.9500 | 0.5510 |
|  |  | 6 | ZOtu782 | 0.8886 | *Lactobacillus* | *Lactobacillus_crispatus* | 0.9300 | 0.3441 |
|  | Comp2 | 1 | ZOtu776 | 0.8734 | *Lactobacillus* | *Lactobacillus_iners* | 1.0000 | 0.8900 |
| CST_IV | Comp1 | 1 | ZOtu304 | 0.9238 | *Prevotella* | *Prevotella_oris* | 1.0000 | 1.0000 |
|  |  | 2 | ZOtu220 | 0.8812 | *Dialister* | *Dialister_invisus* | 1.0000 | 0.9900 |

**Supplementary Table S4. Significant ASVs having differential abundances between CT+ and CT- women.** The 28 ASVs, including two major pathogenic ASVs (ASV sequence #s ZOtu76 and ZOtu95) were identified as differentially abundant between CT+ and CT- (MaAsLin2; unadjusted p-values < 0.05). Among these, 7 *Lactobacillus* ASVs were differentially abundant but had low abundance across 213 women. In contrast, 5 ASVs (ASV sequence #s ZOtu1, ZOtu2, ZOtu11, ZOtu32, and ZOtu374) from *Lactobacillus* *sp.*, which were the most abundant across 213 women but did not exhibit statistically significant differential abundance between CT+ and CT- women, were listed at the bottom of this table. ‘Total read counts’ indicates the number of reads of an ASV across 213 women, and ‘Number of women detected’ shows the number of women who had at least one read count of an ASV among the 213 women. The %ID column shows the identity from BLAST, and the confidence levels for taxonomic assignment at the genus and species levels are indicated in the Confidence level column. Relative abundance was determined using t statistics with indications of whether an ASV was higher in CT+ (Yes) or not (No), which is consistent with the sign of coefficient values. The standard error of the estimated coefficient values is shown in S.E.. **ASVs were assigned to a specific species from among different species that share the same sequence in the V4 region of the 16S rRNA.

< Table is provided in Additional File 2: Supplementary Table S4 >

**Supplementary Table S5. ASVs are predictive of absent CT infection.** These 13 ASVs were identified as predictors for absent CT infection based on random forest classification. The ASVs were detected based on their selection frequency threshold with non-NA feature importance (mean decrease in accuracy), occurring in 9 or 10 out of 10 folds across 100 replications. The 13 ASVs were selected by selection frequencies exceeding 80.

| Relative abundance | Species | ASV | Selection frequency | Total read counts | %ID | Genus | Confidence level | |
| --- | --- | --- | --- | --- | --- | --- | --- | --- |
|  |  |  |  |  |  |  | Genus | Species |
| Higher in CT+ | *Neisseria_gonorrhoeae* | ZOtu95 | 100 | 4,958 | 100 | *Neisseria* | 1.0000 | 0.8900 |
|  | *Prevotella_amnii* | ZOtu829 | 100 | 484 | 95.7 | *Prevotella* | 1.0000 | 0.6900 |
|  | *Syntrophococcus_sucromutans* | ZOtu709 | 100 | 336 | 92.9 | *Syntrophococcus* | 0.3049 | 0.1250 |
|  | *Romboutsia_timonensis* | ZOtu282 | 100 | 213 | 100 | *Romboutsia* | 0.9900 | 0.9603 |
|  | *Syntrophococcus_sucromutans* | ZOtu1814 | 100 | 206 | 92.5 | *Syntrophococcus* | 0.2058 | 0.0658 |
|  | *Syntrophococcus_sucromutans* | ZOtu1481 | 100 | 178 | 91.3 | *Syntrophococcus* | 0.1836 | 0.0404 |
|  | *Trueperella_pyogenes* | ZOtu607 | 97 | 58 | 100 | *Trueperella* | 1.0000 | 0.6800 |
|  | *Rarimicrobium_hominis* | ZOtu470 | 90 | 77 | 100 | *Rarimicrobium* | 1.0000 | 1.0000 |
|  | *Prevotella_brunnea* | ZOtu321 | 88 | 149 | 99.6 | *Prevotella* | 1.0000 | 1.0000 |
|  | *Syntrophococcus_sucromutans* | ZOtu2372 | 82 | 65 | 91.3 | *Syntrophococcus* | 0.2703 | 0.0946 |
| Lower in CT+ | *Lancefieldella_rimae* | ZOtu369 | 100 | 113 | 99.6 | *Lancefieldella* | 0.9900 | 0.9702 |
|  | *Prevotella_oris* | ZOtu304 | 96 | 103 | 100 | *Prevotella* | 1.0000 | 1.0000 |
|  | *Prevotella_melaninogenica* | ZOtu1197 | 95 | 1,481 | 96.8 | *Prevotella* | 1.0000 | 0.4900 |

**Supplementary Table S6. The ASVs consistently chosen for the discrimination between Endo+ and Endo- women by sPLS-DA.** The table lists ASVs that significantly contributed to the first two components of sPLS-DA, discriminating Endo+ and Endo- groups across all women (“All”) and within CST types (“CST I”, “CST III”, and “CST IV”). ASVs with a frequency greater than 0.80 for each component were listed.

< Table is provided in Additional File 3: Supplementary Table S6 >

**Supplementary Table S7. Overview of RDP Classifier 16S trainset No18 raw training database.** When different species share the same V4 region of 16S rRNA reference sequences, 39.7% the V4 reference sequences were redundant for taxonomic assignment to ASVs.

| V4 Reference sequences | Species | Number of V4 Reference sequences | Number of unique V4 Reference sequences | Number of species sharing the same V4 sequence |
| --- | --- | --- | --- | --- |
| 1 | 1 | 10,967 (51.7%) | 10,967 (80.0%) | 1 |
| N | 1 | 1,821 (8.6%) | 786 (5.7%) | 1 |
| 1 | N | 8,407 (39.7%) | 1,133 | 2 |
|  |  |  | 340 | 3 |
|  |  |  | 1,891 | 2 ≤ n ≤ 10 |
|  |  |  | 52 | 10 < n ≤ 20 |
|  |  |  | 7 | 20 < n ≤ 30 |
|  |  |  | 5 | 30 < n ≤ 40 |
|  |  |  | 3 | 40 < n ≤ 50 |
|  |  |  | 0 | 50 < n ≤ 60 |
|  |  |  | 1 | > 60 |
|  |  |  | 1,959 (14.3%) |  |
| Total |  | 21,195 (100%) | 13,712 (100%) |  |

1. Table is provided in Additional File 2: Supplementary Table S4
2. Table is provided in Additional File 3: Supplementary Table S6
